# Metabolism-Informed Bioreactor Model for Transferring Knowledge from Fed-batch to Continuous Biomanufacturing

**DOI:** 10.64898/2026.10.28.762476

**Authors:** Mariana Monteiro, Stacy Bediako, Ricardo Tecson, Gregory Nierode, Pavlos Kotidis, Cleo Kontoravdi

## Abstract

Chinese hamster ovary (CHO) cells remain the dominant platform for monoclonal antibody (mAb) manufacturing, and rising demand is pushing the industry from fed-batch toward continuous operation. Continuous cultures promise sustained high cell densities and volumetric productivities, but their operation depends on reliable real-time monitoring, prediction and control. Limited intracellular observability and sparse extracellular measurements hinder this, leaving a persistent gap between mechanistic understanding and predictive models used in practice.

Here we describe a generalisable, metabolism-informed model that acts as a soft sensor for continuous bioprocess performance, enabling cross-regime deployment. The model uses measured concentrations as inputs and predicts intracellular fluxes. Mass balances at both the intracellular and reactor levels then constrain these fluxes and compute the extracellular metabolite concentrations at the next time point. Because these mass balances hold true across operating modes, they act as a fixed structural prior while the model’s learned dynamics adapt to each regime. Our formulation does not use the predefined rate laws of standard kinetic models, nor does it assume steady state at the reactor level.

We benchmark a genome-scale and a reduced metabolic network, and three optimisation strategies (max ATP, parsimonious FBA, flux sampling), and find that network size affects only flux interpretability, not concentration predictions, while optimisation strategy does not affect flux accuracy. After training on fedbatch data, we compare retraining strategies for continuous operation and find that retraining on limited continuous data works best. This agrees with our measured exchange fluxes, which vary mainly with growth rate rather than operating mode. These results point towards first-principles models that adapt to regime shifts.

## Introduction

Chinese hamster ovary (CHO) cells are the dominant platform for the production of monoclonal antibodies (mAbs), the largest class of recombinant biotherapeutics, due to their capacity to generate human-compatible glycosylation profiles [1, 2]. The growing demand for mAbs has pushed the industry to seek higher productivity and more efficient manufacturing strategies [3], such as operating continuously as it allows the achievement of high cell densities and volumetric productivities [4]. Continuous processing also mitigates the pronounced batch-to-batch variability of fed-batch as cells are held closer to metabolic steady state, with fewer perturbations, for extended periods, supporting more consistent product quality.

Continuous formats such as perfusion are already established in commercial manufacturing for selected products, yet broad adoption across the industry remains limited [5, 6]. Beyond natural risk-aversion and the regulatory hurdles associated with process change, a range of technical and operational challenges constrain wider uptake [7]. During cell line development, knowledge is limited and no metrics exist to assess whether a line is stable enough for continuous operation [8]. Good Manufacturing Practice (GMP) frameworks are only beginning to accommodate continuous mode, standardised process-parameter definitions are still lacking, and filtration is complex while long runs demand costly media volumes [4, 9].

Many of these challenges ultimately reduce to a need to observe and anticipate the state of the culture as it evolves. This is the purpose of process modelling: to infer un-measured intracellular states, forecast extracellular dynamics, and provide the predictive basis for real-time monitoring and control. The maturity of these digital tools, however, remains a barrier. Intracellular mechanisms cannot be directly measured and techniques such as ^13^C flux labelling are costly [10], technically demanding, and ill-suited to routine monitoring. Moreover, many are also restricted to metabolism, leaving functions like protein synthesis and secretion poorly characterised. Extracellularly, key variables such as metabolite concentrations are measured infrequently, as sampling is overly manual [11] or online measurement prohibitively expensive. Thus, models have attempted to approximate intracellular structure with semi-empirical kinetic expressions (such as Monod saturation kinetics) fitted to extracellular data [12–15], but since these are calibrated under specific operating conditions, they inherit regime-specific knowledge that limits transferability across modes of operation.

Hybrid modelling approaches have sought to bridge the gap between data observability on the extracellular scale and unobservable mechanisms on the intracellular scale. These modelling approaches typically embed neural networks within predefined kinetic reactor models [16, 17] or augment mass balances with data-driven corrections [18], and have been extended to incorporate uncertainty quantification [19]. These approaches still keep the mechanistic core kinetically parameterised. Alternatively, [20] has proposed a relaxation to the mass balance at reactor level as a soft residual constraint in a physics-informed architecture, where the rates are learned from a feed-forward neural network. Parallel efforts have integrated constraint-based metabolic models with statistical or learning-based layers to refine intracellular flux estimates by restricting feasible solution spaces [21–23]. More recently, [24] employed a hybrid formulation to predict specific reaction rates that served as constraints for a genome-scale model, which was later uniformly sampled. Many of these hybrid approaches share the limitation of the mechanistic models discussed above, as their assumptions are tied to specific operating conditions, compromising transferability under regime shifts.

To overcome the regime-specificity of existing models, we develop a metabolism-informed model that acts as a soft sensor for bioprocess performance, embedding reactor and intracellular mass balances into a recurrent learning architecture without assuming predefined kinetic laws. The model takes measured extracellular metabolite concentrations as input, predicts intracellular fluxes, and passes these through the embedded mass balances to compute the next time point’s concentrations. We separate structural priors – stoichiometry and reactor-level mass balance, which are determined by chemistry and therefore invariant across process conditions and modes of operation [25] – from kinetic priors, which are learned from data and permitted to adapt across regimes. Steady state is not assumed at the reactor level, and during training the predicted fluxes are encouraged to lie within intracellular bounds supplied by NEXT-FBA [23], a hybrid stoichiometric/data-driven approach that infers physiologically informed bounds from measured exchange fluxes and ^13^C labelling data, providing a structured prior on the feasible flux space with-out requiring direct intracellular measurements. We evaluate the model on the transfer from fed-batch to continuous mAb production with limited continuous culture data.

Trained exclusively on six industrial fed-batch cultures, the metabolism-informed model predicts metabolite concentrations and exchange fluxes with comparable accuracy to a fully data-driven baseline. We evaluate the transferability capacity of the fed-batch-trained model to continuous operation, and find that fully fine-tuning the model parameters yields the best performance. Predicted fluxes are shown to be robust to the choice of metabolic optimisation strategy (max ATP, parsimonious FBA, flux sampling) and metabolic network size (genome-scale versus reduced-scale), though network size affects the interpretability of the recovered fluxes. Growth rate rather than operating mode emerges as the dominant axis of variation in the measured exchange fluxes. Across model configurations, the predicted intracellular fluxes suggest regime-shift signals that can inform experimental design for continuous culture characterisation. These results point towards digital models grounded in first-principles knowledge that can adapt to regime shifts.

## Methodology

### Experimental data

Nine CHO cell culture production runs were analysed, comprising six fed-batch (FB1-FB6) and three perfusion (C1-C3) experiments. Cultures were performed in 3 L stirred tank reactors using a proprietary platform process for recombinant mAb production.

Viable cell density and viability were measured using a Beckman Coulter ViCell XR. mAb titre, glucose (Glc), lactate (Lac), ammonia (Amm), glutamine (Gln), and glutamate (Glu) were measured using a Roche CEDEX. The concentrations of amino acids alanine (Ala), asparagine (Asn), aspartate (Asp), glycine (Gly) and serine (Ser) were measured using a Repligen REBEL XT Analyzer. Measurement error values were available for amino acids, provided by the instrument vendor. For the remainder, a nominal value of 10% was assumed. Missing time-series measurements were interpolated using cubic splines.

Fed-batch cultures were initiated at 1.6 L and operated at 35°C, 50% dissolved oxygen and 350 rpm agitation. Nutrient feed was added to the reactor every 2-3 days, starting on day 3, for a total of 20% of the initial working volume. Glc was supplemented up to 5 g/L if the daily measured Glc was below 4 g/L, using a 400 g/L stock solution.

For the perfusion experiments, the inoculation volume was set at 2 L, and perfusion of proprietary cell culture media started on day 1, with bleeding on day 2 to maintain a target viable cell concentration of 30 × 10^6^ cells*/*mL or lower. The harvest rate was back-calculated from the reactor volume, media addition, and bleed rates. This adjustment was necessary because the measured and calculated volumes did not match, and the harvest rate was deemed less reliable, although error in the bleed rate cannot be excluded.

### Model Structure

Inputs to the model consist of measured metabolite concentrations, associated measurement errors, decision variables (including reactor feeding composition and bleed and filtrate volumes) and bioreactor volume. With the exception of the measurement errors which are appended directly to the concatenated encoder, these inputs are processed through three parallel long short-term memory modules, each encoding the temporal dynamics of one input group. The resulting latent representations are concatenated and mapped to two output heads: a metabolic model head, predicting a flux vector, and a death-rate head. The concentrations of metabolites at the reactor level are obtained by integrating the bioreactor mass balance forward over one measurement interval. This integration has as inputs the predicted fluxes from the metabolic head and decision variables (i.e., feeding regime).

#### Integration of the Reactor Mass Balance

Rather than predicting concentrations directly, the model advances the measured state at *t* to a prediction at *t* + 1 by integrating the bioreactor mass balance over the interval, using the exchange fluxes and the biomass flux read out of **v** and the death rate *k*_*d*_ from its own head. Cells are removed by the bleed only and metabolites by bleed and filtrate together. Both balances are integrated analytically over the interval:

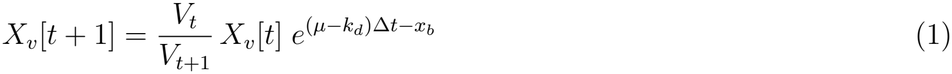

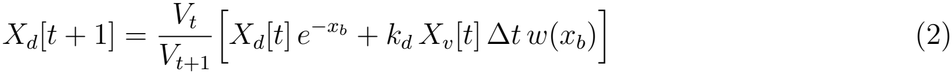

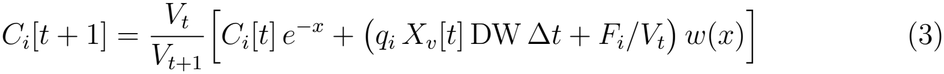

Here *V*_*t*_ and *V*_*t*+1_ are the measured volumes; *µ* is the biomass flux and *q*_*i*_ the exchange flux of species *i*, both components of **v**; *F*_*i*_ is the amount of species *i* supplied by the feed and media streams over the interval; *x* = (Filtrate + Bleed)*/V*_*t*_ and *x*_*b*_ = Bleed*/V*_*t*_ are the exchanged volume fractions; *w*(*x*) = (1 − *e*^−*x*^)*/x* accounts for the dilution of material added at a constant rate over the interval; and DW is the dry weight per cell. The left endpoint of every interval is the measured state, during training and at evaluation, so all reported predictions are one-step-ahead.

#### Composite Loss

Model training is guided by a composite objective (loss function) that balances predictive accuracy with metabolic consistency. The loss function is composed of the following terms, which will be explained next:

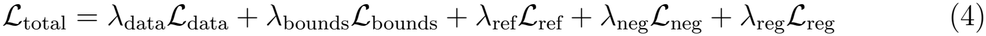

#### Data Loss

The data term penalises discrepancies between predicted and measured concentrations, where the prediction is the one-interval integration above from the measured state at *t* to the measurement at *t* + 1.

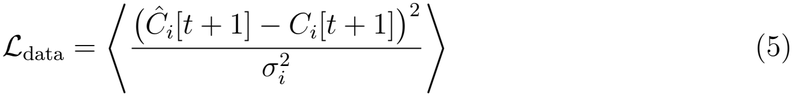

#### Metabolic Loss

The metabolic head produces a full intracellular flux vector as **v** = *Nz*, where N is the null-space basis of the stoichiometric matrix S, and z are coordinates learned by the network. Because *SN* = 0 (by definition of the null space), stoichiometric balance *S***v** = 0 holds exactly at every training and inference step. The network’s *Nz* term cannot violate stoichiometric balance but can adjust **v** anywhere in the null space, including the reactions corresponding to measured exchange channels, which is how the exchange fluxes driving the mass balance are produced.

Two loss terms carry metabolic prior knowledge onto **v**, ℒ_*bounds*_ and ℒ_*ref*_, neither of which enters the forward pass. Intracellular bounds are supplied by NEXT-FBA [23], which receives measured exchange fluxes and returns intracellular bounds informed by ^13^*C* labelling data. These are combined per time point with reversibility from the network file, ±*kσ* windows on the measured exchange fluxes (*k* = 2, *σ* from the reported measurement error), and a one-sided bound on growth: the measured net growth rate *µ* − *k*_*d*_ is a lower bound on *µ*, not an estimate of it, so no upper window is derived from it. The first term is a soft quadratic bounds penalty:

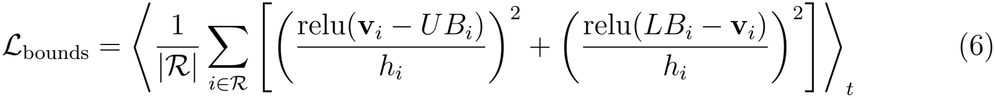

where ℛ is the set of reactions with finite bounds, *LB*_*i*_ and *UB*_*i*_ are the per-time-point bounds, and *h*_*i*_ is the training-set median half-width of reaction *i*’s window, used to non-dimensionalise the penalty. A violation is therefore scored relative to how much room the reaction typically has.

The second term pulls the reactions unconstrained by data toward a reference flux distribution **v**_ref_ computed offline for every time point. Three solution strategies are available for computing **v**_ref_ : parsimonious flux balance analysis (pFBA) [26], maximisation of ATP, and flux sampling [27]. These are described in more detail in the Supplementary Information. All three optimise inside the same polytope, defined by *S***v** = 0 and the bound windows above. The reference is obtained in two stages per time point: (i) a linear programme first finds the minimal weighted relaxation of the windows that makes the polytope non-empty (measurement windows are weighted 100× more heavily than NEXT-FBA intervals, and reversibility is never relaxed); and (ii) the objective is then optimised subject to that relaxation.

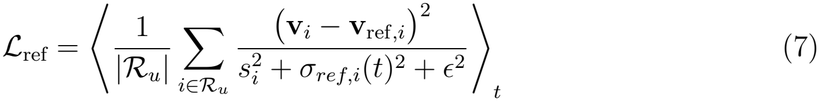

where ℛ_*u*_ excludes the measured exchange channels and biomass (including them would duplicate the data term and make solver agreement on those channels true by construction), *s*_*i*_ is the spread of the reference over the training time points, and *σ*_*ref,i*_(*t*) is the sampling spread (zero for the point solvers), and *ϵ* is a small constant that keeps the denominator non-zero. For sampling, the additional term down-weights reactions with a wide spread across samples, so the reference pulls mostly on reactions that are well constrained.

Figure 3 provides a detailed account of the architecture behind the metabolic loss structure. In phase 0, three inputs are specified: the metabolic network model (CHOm-pact [28] or iCHO2441 [29]), experimental data and the metabolic optimisation strategy (pFBA, max ATP or flux sampling). In phase 1, three elements are precomputed and fixed: null-space basis of the stoichiometric matrix, the flux bounds (coming from either NEXT-FBA [23] or from experimental data) and the reference flux (computed by the chosen optimisation strategy within these bounds). In phase 2, executed at every iteration during training, the following occurs: the metabolic head predicts the independent fluxes (corresponding to the degrees of freedom left by stoichiometry); the intracellular mass balances reconstruct the full flux vector from them so that the steady-state assumption holds by construction; and the metabolic loss penalises the violations of the precomputed bounds and pulls the reactions towards the reference.

**Figure 1:**
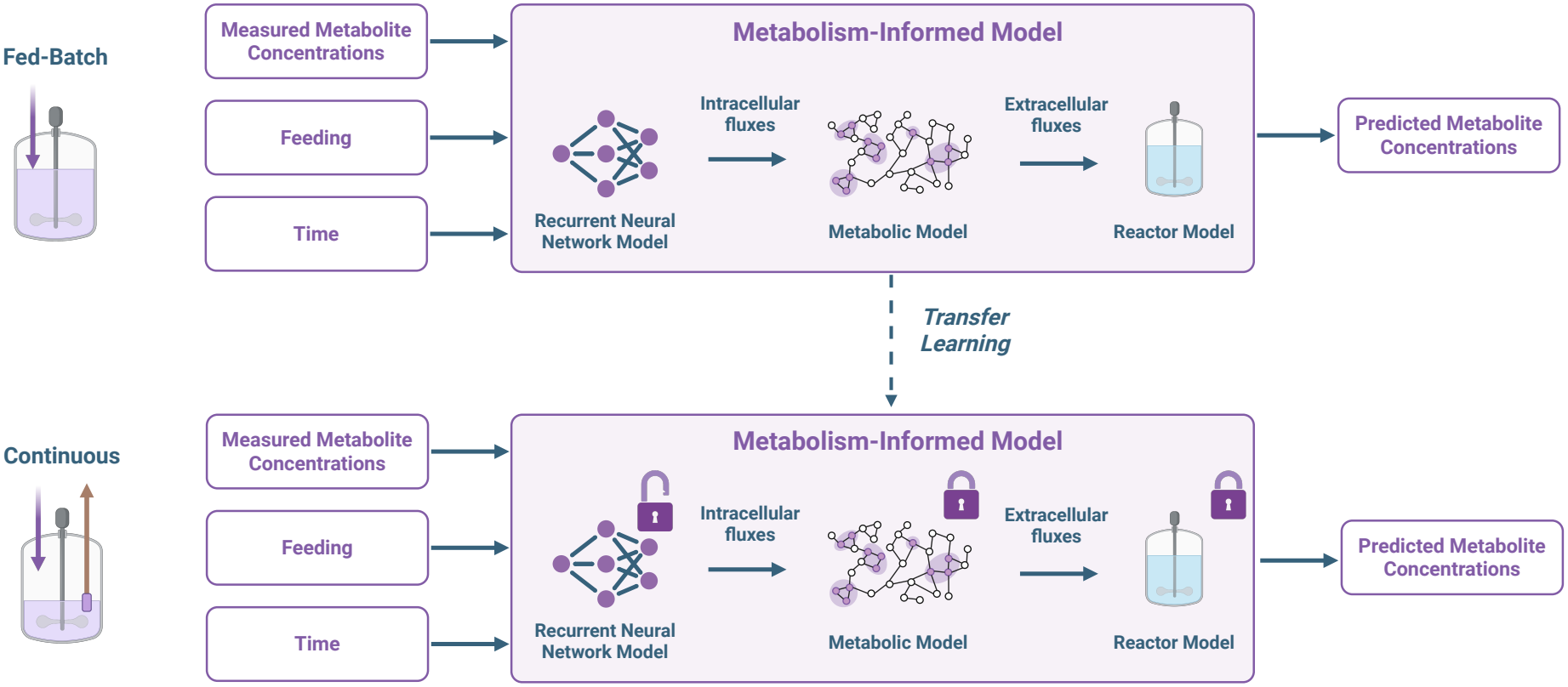
Graphical Abstract. Proposed architecture of the metabolism-informed model. A recurrent neural network predicts intracellular fluxes from measured concentrations, which pass through metabolic and reactor models enforcing mass balance to predict the next concentrations. Trained on fed-batch data, the model is then adapted to continuous operation via transfer learning: the recurrent neural network is retrained on continuous data, while the metabolic and reactor models remain fixed. Image created in BioRender.

**Figure 2:**
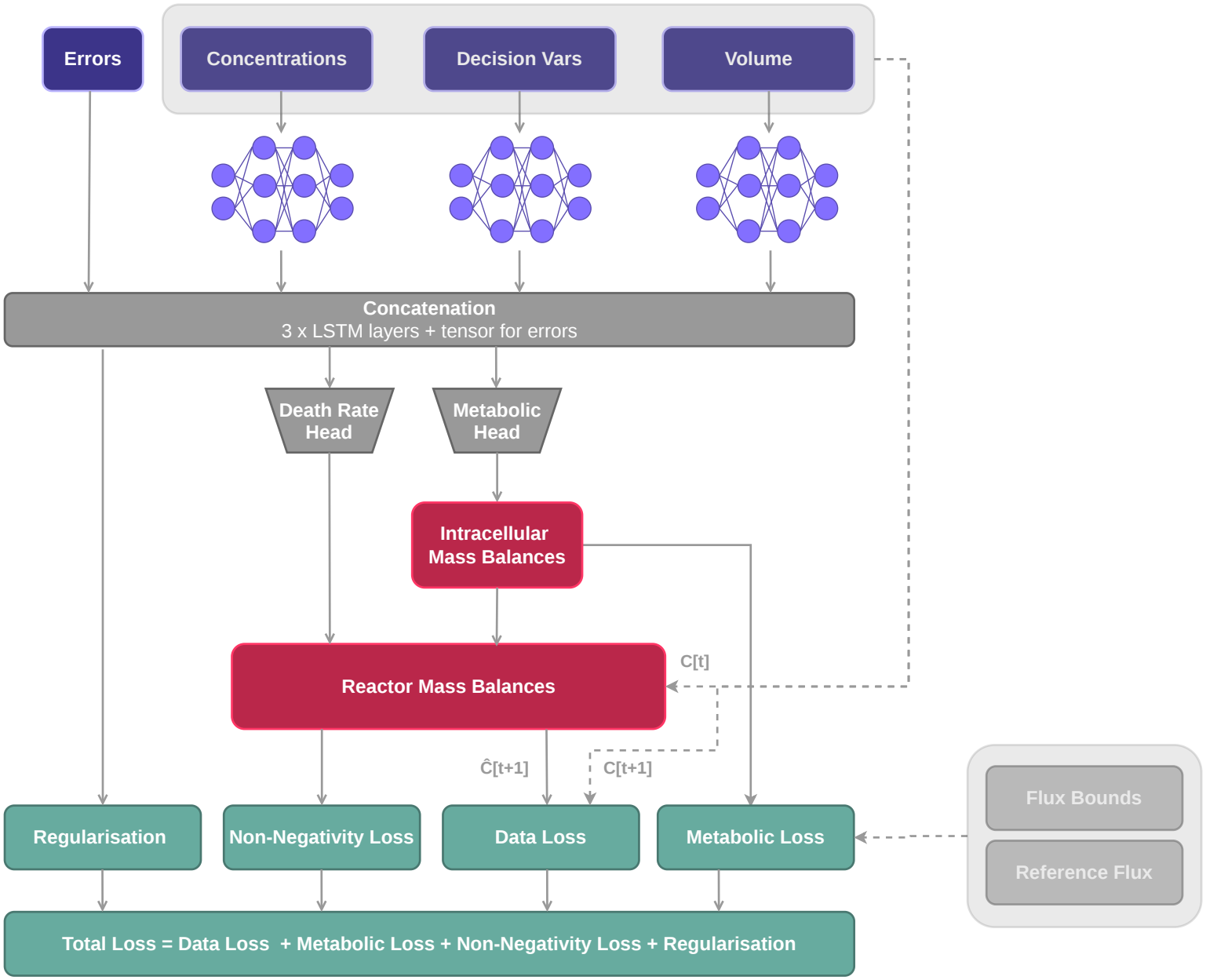
Metabolism-informed model structure. Metabolite concentrations, their respective measurement errors, decision variables (feed, media and bleed rates) and reactor volume are the inputs to the metabolism-informed model. With the exception of the measurement errors, each set of inputs is fed to a long short-term memory (LSTM) encoder, and then concatenated back into a shared representation. This is then mapped onto a metabolic head and a death-rate head. The former predicts a set of independent fluxes, which are reconstructed through the null space of the stoichiometric matrix into a full flux vector that satisfies the intracellular mass balances. Both these fluxes and the death rate are introduced to the reactor mass balances module, which integrates the measured state at time *t* forward over one measurement interval, using the measured volumes and feed, media and bleed information to give a set of predicted concentrations at *t* + 1. These enter the Data Loss module, which computes their difference from the measured concentrations at *t* + 1. The Metabolic Loss module penalises predicted fluxes that fall outside the precomputed bounds and pulls the unmeasured reactions toward the precomputed reference flux. The non-negativity loss penalises predicted concentrations that cross zero, and an ℓ_2_ regularisation on the network weights softly penalises overfitting. The total loss is a weighted sum of the four terms, with the Metabolic Loss comprising the bounds and reference terms. All predictions are one-step-ahead from the measured state.

**Figure 3:**
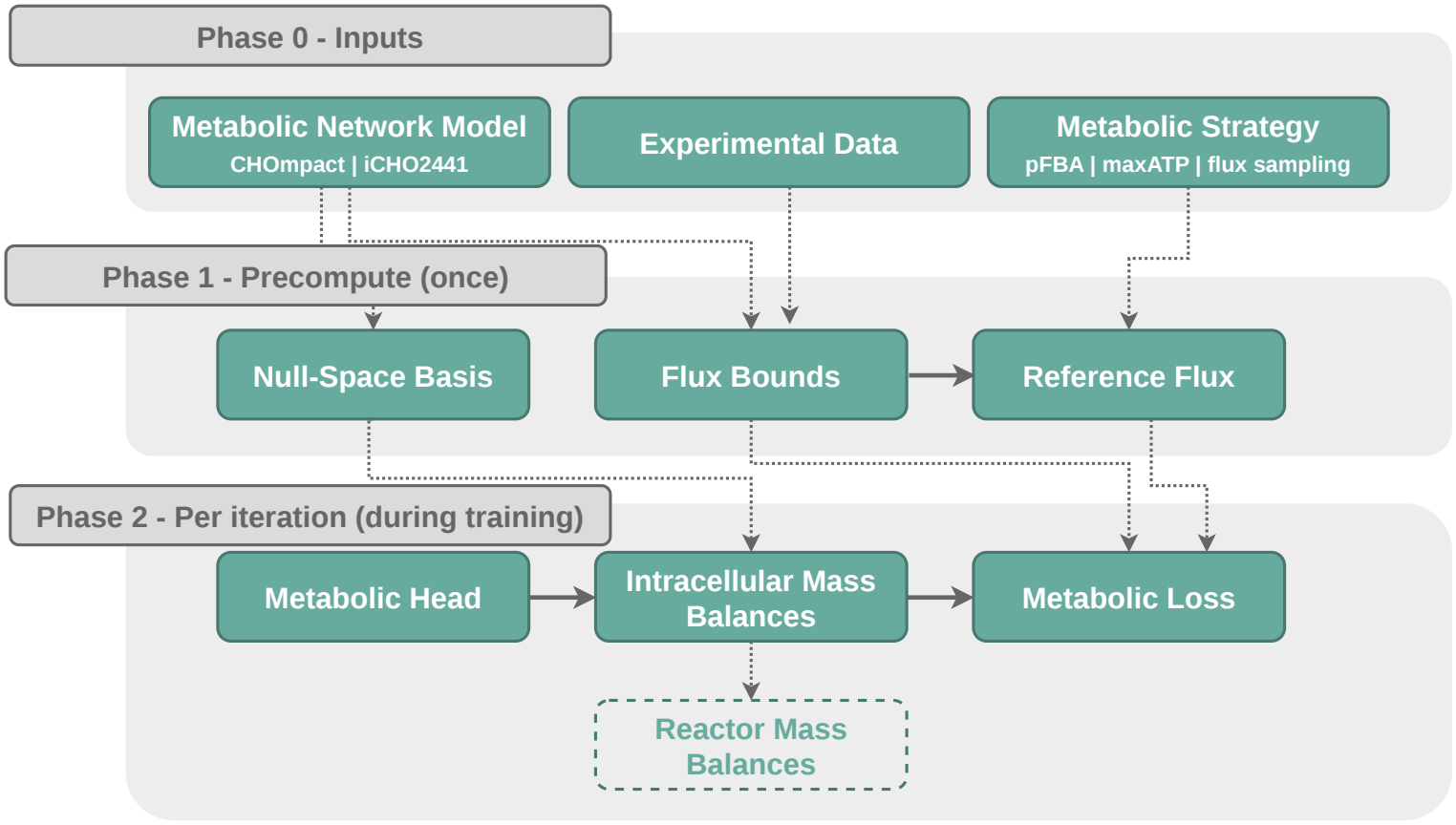
Three-phase architecture of the metabolic loss pipeline.

The decision to implement a null-space parameterisation was primarily motivated by the need to reduce the network’s output dimensionality. Predicting a full intracellular flux vector would significantly increase the number of parameters in the final layers (especially for larger metabolic models). Instead, the network predicts only the null-space coordinates *z*, and the full flux vector is obtained as **v** = *Nz*. A useful consequence is that stoichiometric balance *Sv* = 0 becomes a hard architectural constraint rather than a soft loss penalty. The dimension of *z* is that of the null space of *S*, so it grows with the network.

#### Non-Negativity Loss and Regularisation

The non-negativity loss is formulated as follows:

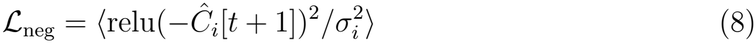

It penalises integrated concentrations that cross zero. Given that the consumption term in the mass balance is a zero-order sink, an over-predicted uptake can remove more of a species than is present in one interval. To avoid overfitting, ℒ_reg_ is defined as the squared ℓ_2_ norm of the parameters.

At inference, the network produces flux predictions through the same forward pass: the LSTM hidden state is mapped to *z*, and the full flux vector is reconstructed as described previously. However, because ℒ_bounds_ is a soft penalty applied only during training, the predicted flux is not guaranteed to lie within the bound-feasible polytope at inference. In particular, the bounds penalty is scaled by the width of each reaction’s window (*h*_*i*_). Irreversible reactions with a loose upper bound (e.g. 1000) therefore have a very wide window, so a negative value on such a reaction is barely penalised during training and can appear in the predicted flux distribution. We quantify this in the Results Section and discuss its implications for the regime-shift analysis.

### Step 0: Hyperparameter Tuning

The hyperparameters of the metabolism-informed model, including the contribution of each loss to the total loss, were tuned using the Optuna package [30]. The search covers the loss weights *λ*_bounds_, *λ*_ref_ and *λ*_neg_ (log-uniform), the ℓ_2_ weight, the learning rate, the LSTM width, the dropout before the heads, and the length of the metabolic-loss warm-up; the number of LSTM layers is fixed at one. *λ*_data_ is fixed at 1.

200 trials were run, each training the model on two fixed fed-batch folds for at most 150 epochs, with an early stopping patience of 30 epochs; the objective is the mean over the two folds of the best validation data loss, and unpromising trials are pruned after their first fold. Trials are ranked on the data loss only, because the metabolic bounds are built from the held-out experiment’s own measurements. Hyperparameters are tuned once, on two of the six folds, and then applied to all six folds of the leave-one-out training described next. The data-driven baseline is tuned by the same procedure over its own hyperparameters (width, ℓ_2_, learning rate, *λ*_neg_, dropout). The Adam optimiser was used throughout.

#### Step 1: Train Fed-Batch Model

The model is first trained on fed-batch data, using a leave-one-out cross-validation scheme, over the six experiments. The metabolic loss weights are linearly warmed up from 0 to their target weight over the first epochs (the warm-up length is also a hyperparameter) to avoid early instability during training. There is an early stopping mechanism that monitors the validation data loss and restores the best weights if there is no improvement after 75 epochs, within a budget of 1000 epochs. For subsequent use (Section Step 2: Fine-Tune Continuous Model), a consensus model is trained with all fed-batch experiments, for better data usage.

#### Step 2: Fine-Tune Continuous Model

The consensus model from Step 1 (Section Step 1: Train Fed-Batch Model) is fine-tuned with less abundant continuous data, also using a leave-one-out cross-validation scheme, over the three experiments. Similarly to before, there is a warm-up scheduler for the losses and an early stopping mechanism. The learning rate is halved from Step 1. The decision-variable encoder has the same six inputs in both modes (feeds, filtrate, bleed and media), with streams absent from a mode set to zero. As filtrate, bleed and media are zero throughout fed-batch training, their input weights stay at their initial values in the consensus model, and are only trained in transfer learning (TL) strategies T1, T3 and T4 (see below).

Five fine-tuning strategies were benchmarked, testing different hypotheses about the domain gap. The trainable parameters partition exactly into the three LSTM encoders and the two output heads (the null-space flux head and the death-rate head), so a strategy is defined by which of the two blocks is trained. Table 1 summarises the main differences.

**Table 1:** TL strategies. Each strategy fine-tunes a different subset of the model on continuous-culture data, using the fed-batch-trained consensus model as initialisation.

| **Strategy** | **Hypothesis** | **Encoders**<br>(3 LSTM) | **Heads**<br>(flux, death) |
|---|---|---|---|
| T0: no fine-tune | Fed-batch model applied directly | - | - |
| T1: full fine-tune | All components retrained | train | train |
| T2: encoders frozen | Temporal representation transfers; the state→flux map shifts | freeze | train |
| T3: heads frozen | State→flux map trans-fers; the dynamics shift | train | freeze |
| T4: from scratch | No fed-batch initialisa-tion | train | train |

In the full fine-tuning (T1) case, every parameter in the model is retuned to adapt to the continuous regime, and the fed-batch model parameters represent initial guesses.

In the case of T2, the three LSTM encoders are frozen, so the temporal representations learned from fed-batch mode are preserved, and the two heads relearn how to map those fixed representations onto continuous-culture fluxes and death rate. In the T3 strategy, the heads are frozen and only the encoders adapt, testing whether the representation-to-flux map learned in fed-batch holds and it is the temporal dynamics that differ. The death-rate head is frozen together with the flux head in T3 so that T2 and T3 remain an exact partition. T4 re-initialises every weight, so it shares the architecture and hyperparameters but no fed-batch knowledge. Finally, strategy T0 applies the consensus model with no weight adaptation, with only the input scaling refitted to the continuous data. Together the five strategies bracket the value of the fed-batch initialisation, from the fed-batch model applied without any continuous data (T0) to a model trained on continuous data alone, with no fed-batch initialisation (T4).

## Results

### Part I: Model performance

Throughout this section, the metabolism-informed model uses the CHOmpact metabolic network and parsimonious FBA (pFBA) as its solution strategy to resolve the metabolic mass balances (both decisions are benchmarked in Part II). As described in the previous Section, we first tuned the hyperparameters of the metabolism-informed model and of its fully data-driven benchmark. Table 2 reports the search ranges and the tuned values for each.

**Table 2:**
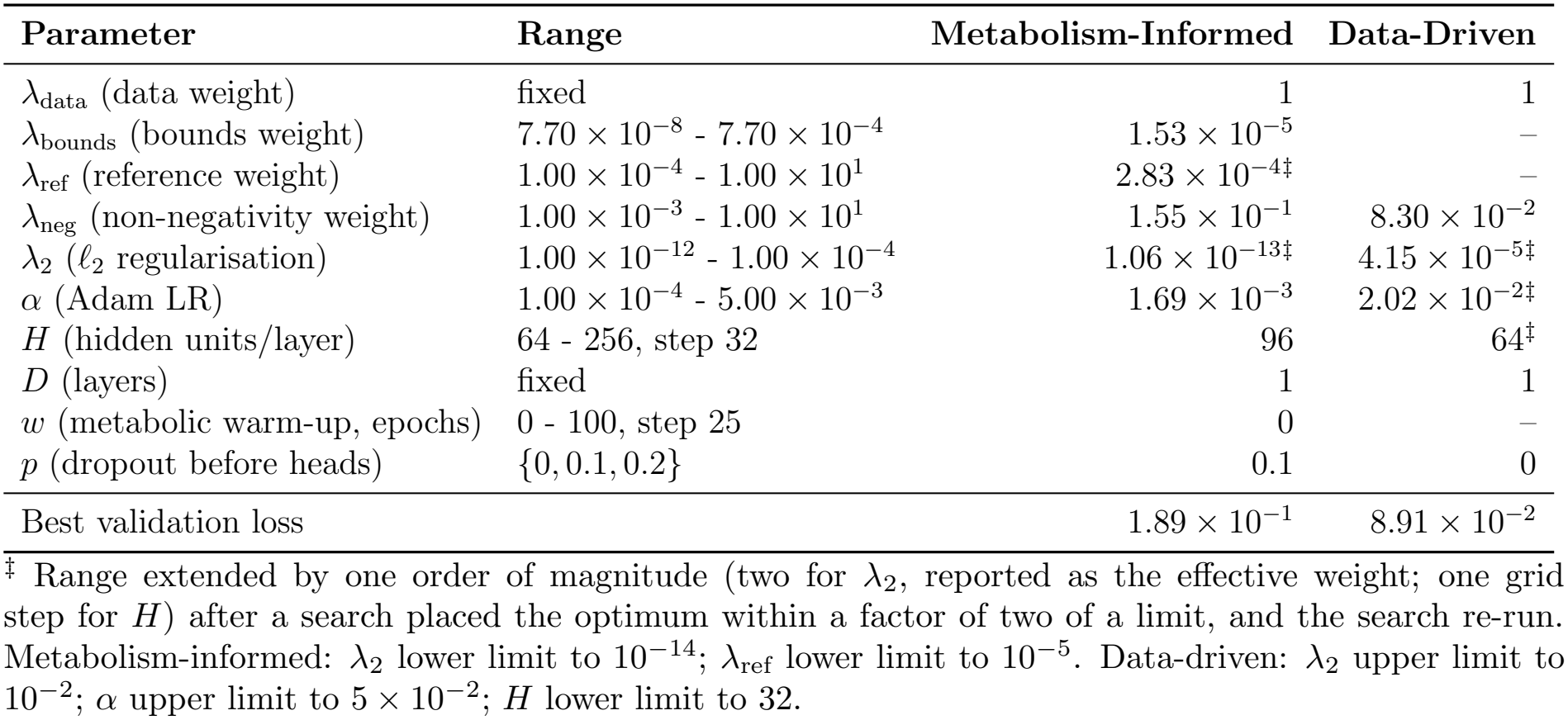
Hyperparameter search ranges and optimised values, for the metabolism-informed and data-driven models. Ranges are log-uniform unless noted. Both searches started from these ranges, and values marked ^*‡*^ were found after the range was extended (see note). The metabolic weights and warm-up do not apply to the data-driven model, which has no metabolic terms. Best validation loss is the mean over the two hyperparameter tuning folds of the best validation data loss (200 trials in the final search for each model).

The number of layers was fixed at 1 in both models. In an earlier version of the model (results not shown), the search chose a single layer for both model types, suggesting that the temporal dynamics are simple enough that the data do not support deeper architectures; increasing the depth of an LSTM would also substantially increase the number of parameters for no gain in the information available to fit them.

The tuned settings suggest that the optimal structure for the data-driven model is relatively reduced in size (64 units), damped (*λ*_2_ eight orders of magnitude above the metabolism-informed model), that learns fast (learning rate of Adam one order of magnitude above). On the other hand, the metabolism-informed model is wider, learns more slowly and is effectively unregularised by ℓ_2_, with dropout as its only explicit regulariser. One possible explanation is that the mechanistic priors already constrain the model, although the search did not test this directly. The warm-up of the metabolic losses was tuned to 0, which suggests that their weight in the overall loss is small enough not to benefit from one.

The non-negativity weight is higher in the metabolism-informed model, which points to its necessity in the model despite the existence of priors. In Part II, we discuss how the data contributes to negative predicted concentrations.

The best validation loss of the data-driven model is about half that of the metabolism-informed model. This suggests that the metabolism-informed model does not achieve a better concentration prediction than the data-driven model, for the present data. Given that the validation loss only accounts for concentration (and not flux) data, this is expected. The priors of the metabolism-informed model act on the intracellular fluxes, which the concentration data do not constrain.

In fed-batch mode, the two model structures achieve comparable accuracy in both concentration and exchange flux prediction (Fig. 4). It is important to note that the data-driven model does not predict fluxes; instead these are inverted through mass balance for validation purposes.

**Figure 4:**
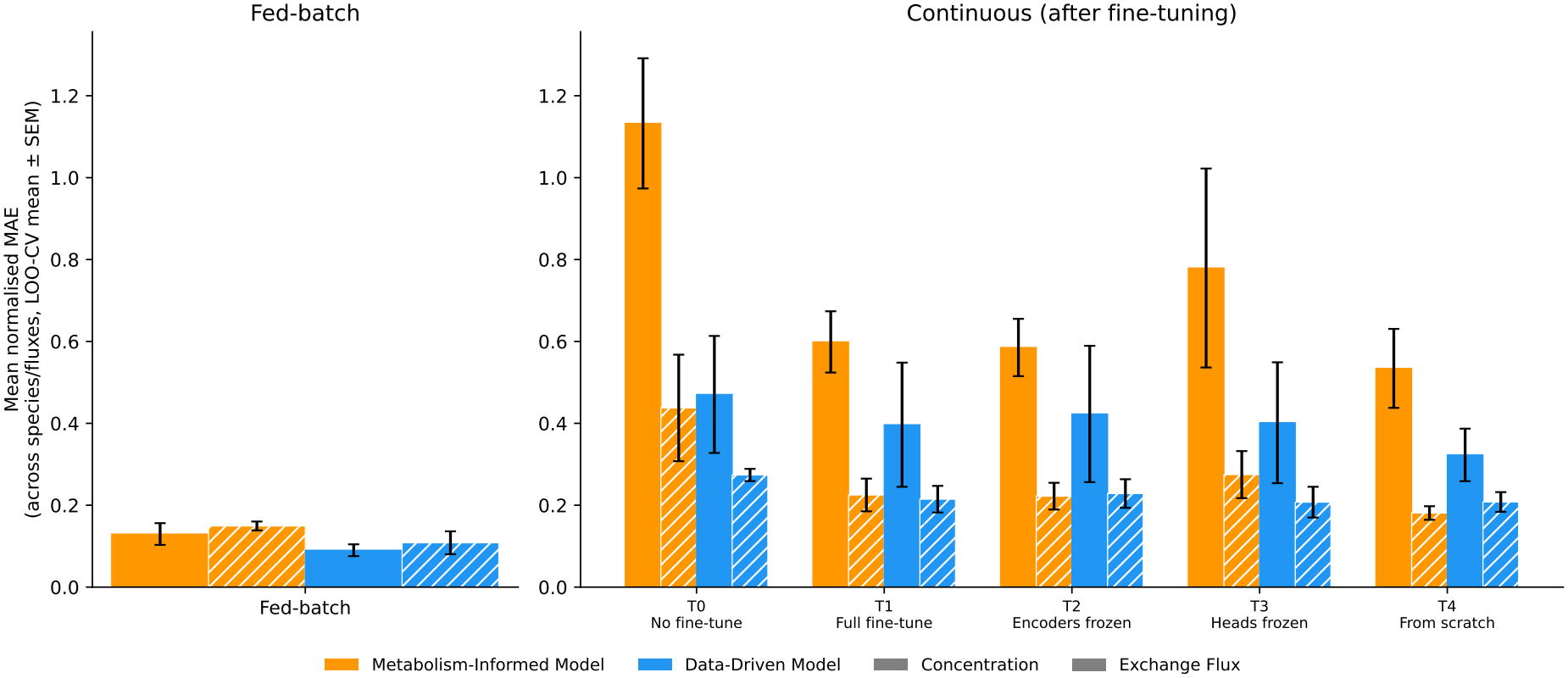
Normalised mean absolute error (MAE) for concentration and exchange fluxes across all species, averaged over all held-out folds by metabolism-informed model against a fully data-driven model. Left-hand side pertains to fed-batch mode and right-hand side continuous mode, after different types of fine-tuning.

In continuous mode, the two models diverge, with the data-driven model holding an edge in both concentration and flux prediction. This regime favours this advantage, as the continuous training set contains only three experiments of similar process conditions, under which a lower-variance data-driven fit is preferred at the cost of higher bias. The advantage is most noticeable in transfer learning strategies T0 (no fine-tuning, meaning using the fed-batch model directly on continuous data, without retraining) and T3 (metabolic and death rate heads frozen). In the remaining transfer learning strategies, the differences are within variance.

### Part II: Evaluation of model generalisation

Tuning the individual loss weights (Part I) tends to favour the data term, allowing relaxation in the mechanistic terms. In Part II we instead weight each term contributing to the loss equally. Using a model trained with *λ*_*ref*_ = 0 (so all three solvers give the same training signal), the unweighted loss terms were evaluated per solver and used to set *λ*_*bounds*_ = ℒ_*data*_*/*ℒ_*bounds*_ and *λ*_*ref*_ = ℒ_*data*_*/*ℒ_*ref*_, so each term matches ℒ_*data*_ in magnitude for that model. The calibrated *λ*_*ref*_ values were similar across solvers, so the pFBA value was used for all three. The remaining hyperparameters are fixed and identical across configurations (hidden size 96, 1 layer, dropout 0.1 and batch size 4). This weighting is used for all results in Part II, including the comparison of fed-batch and continuous operation in the sections that follow.

Flux sampling was not run on iCHO2441 because building the sampling reference is computationally prohibitive at that network size: a single time point took 39 min end to end (dominated by flux variability analysis over 12,674 linear programmes on 6337 reactions), which extrapolates to roughly 54 h per training set and over 500 h for the full fed-batch and continuous grid, roughly 100 times the compute used for all other Part II runs combined. This section thus compares three solvers on CHOmpact and two (pFBA and max ATP) on iCHO2441.

#### Equal loss weighting preserves predictive accuracy

Under leave-one-out cross-validation, the metabolism-informed model accurately predicts most species, within their measurement errors (see Figure 5 for CHOmpact’s case and Supplementary Figure 14 for iCHO2441). Per-fold breakdowns of concentration prediction for each model can be found in Supplementary Figure 16.

**Figure 5:**
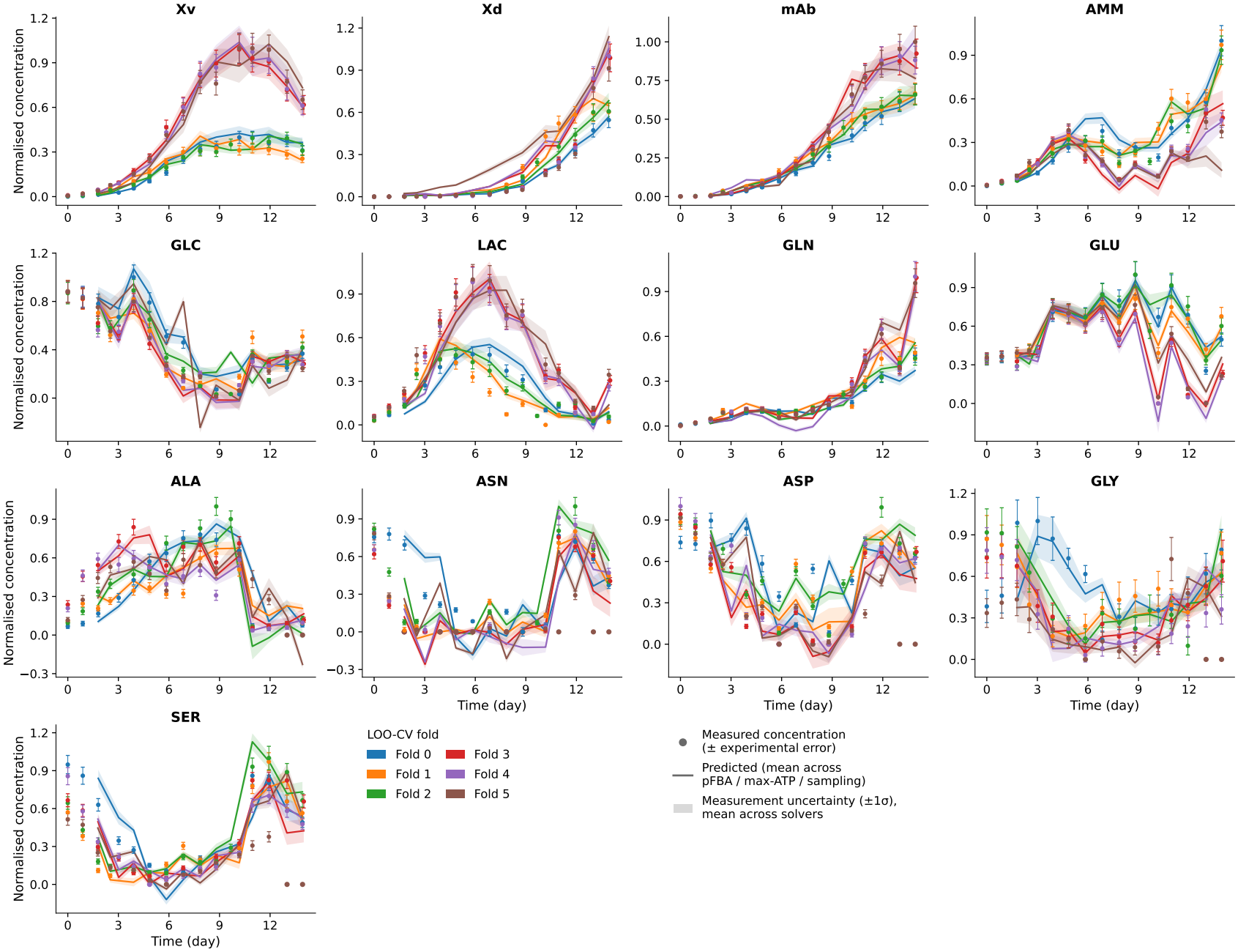
Concentration trajectory predictions for cross-validation experiments in fedbatch mode (model = CHOmpact). Each solid line is the mean prediction across pFBA, max ATP, and flux sampling.

mAb and viable cell density (normalised MAE of around 0.1) are tracked best, with the remaining metabolites below 0.3. The exceptions are asparagine, aspartate and serine in fold 5, reaching almost 0.5. This is observed consistently across model sizes and optimisation strategies, suggesting this discrepancy is not model structure related.

Exchange fluxes (Figure 6 for CHOmpact and Supplementary Figure 15 for iCHO2441) are predicted with the same range of accuracy as concentration for most metabolites (typically 0.10–0.30, see Supplementary Figure 17). The exceptions sit at the lower and higher ends of the spectrum of magnitude of fluxes. The largest errors fall on mAb at the low end of the measured range. In the case of alanine, asparagine, glucose, lactate and serine, the model systematically underpredicts the higher flux values. This suggests a regression towards the centre of the training distribution: at the sparsely sampled high-flux tail, the network learns a feasible distribution that underestimates the true magnitudes.

**Figure 6:**
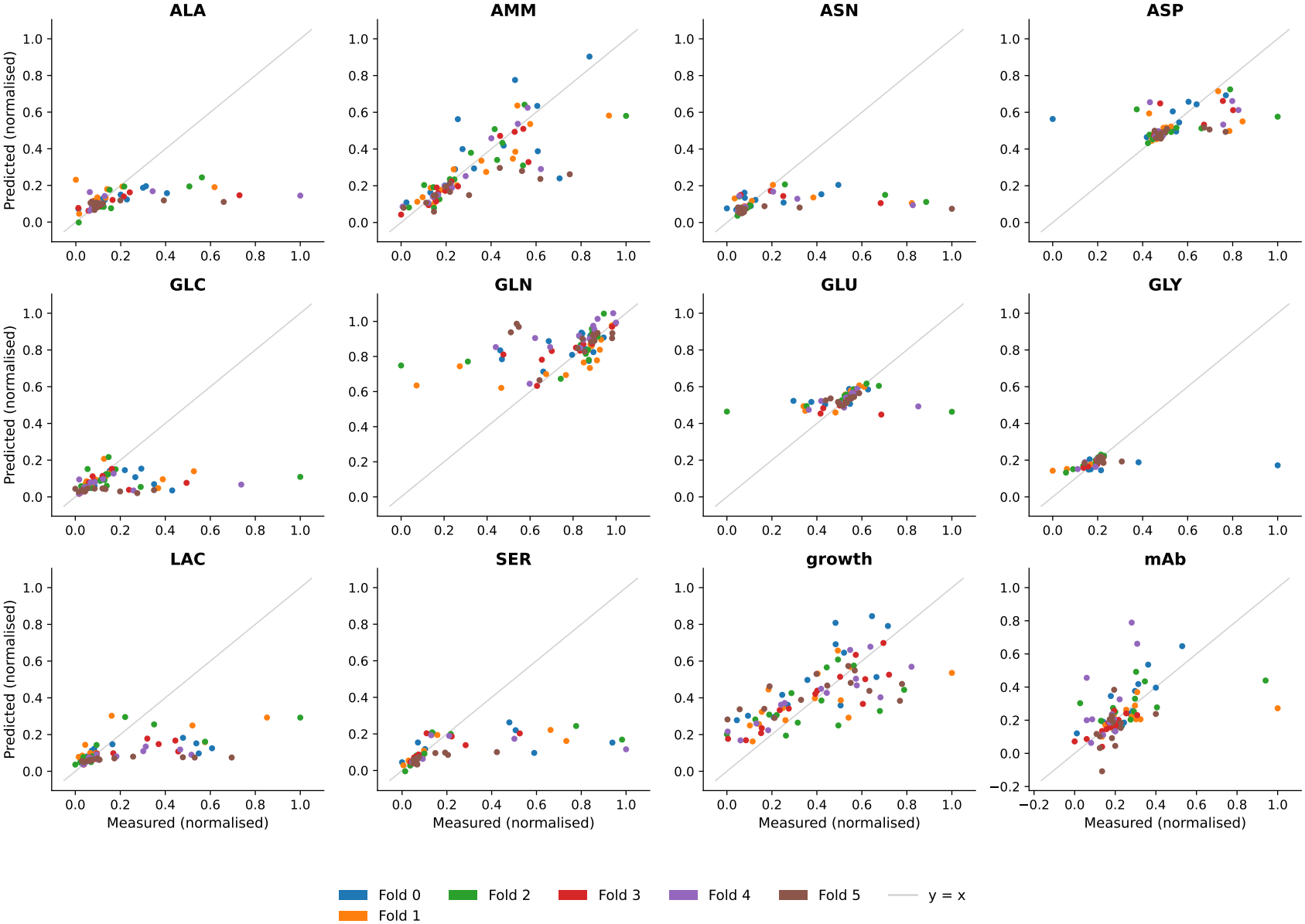
Exchange flux predictions for cross-validation experiments in fed-batch mode (model = CHOmpact).

The results show that the predictive accuracy of the model outputs is robust to the choice of metabolic optimisation strategy. This is positive, as it means that the optimal choice of objective function is less of a concern with this model structure. This makes the proposed model an appropriate option when there is no agreement on which optimisation strategy to use. Using a genome-scale model does not seem to offer an advantage on predictions, when compared to the reduced-size network.

#### Genome-scale resolution expands the flux space without improving concentration predictions

CHOmpact (144 reactions) and iCHO2441 (6337 reactions) achieve indistinguishable permetabolite normalised MAE across all 13 measured species. Figure 7 shows the improvement (in %) of iCHO2441 over the reduced-size model CHOmpact for exchange flux prediction, where positive values indicate metabolites that are better described by iCHO2441.

**Figure 7:**
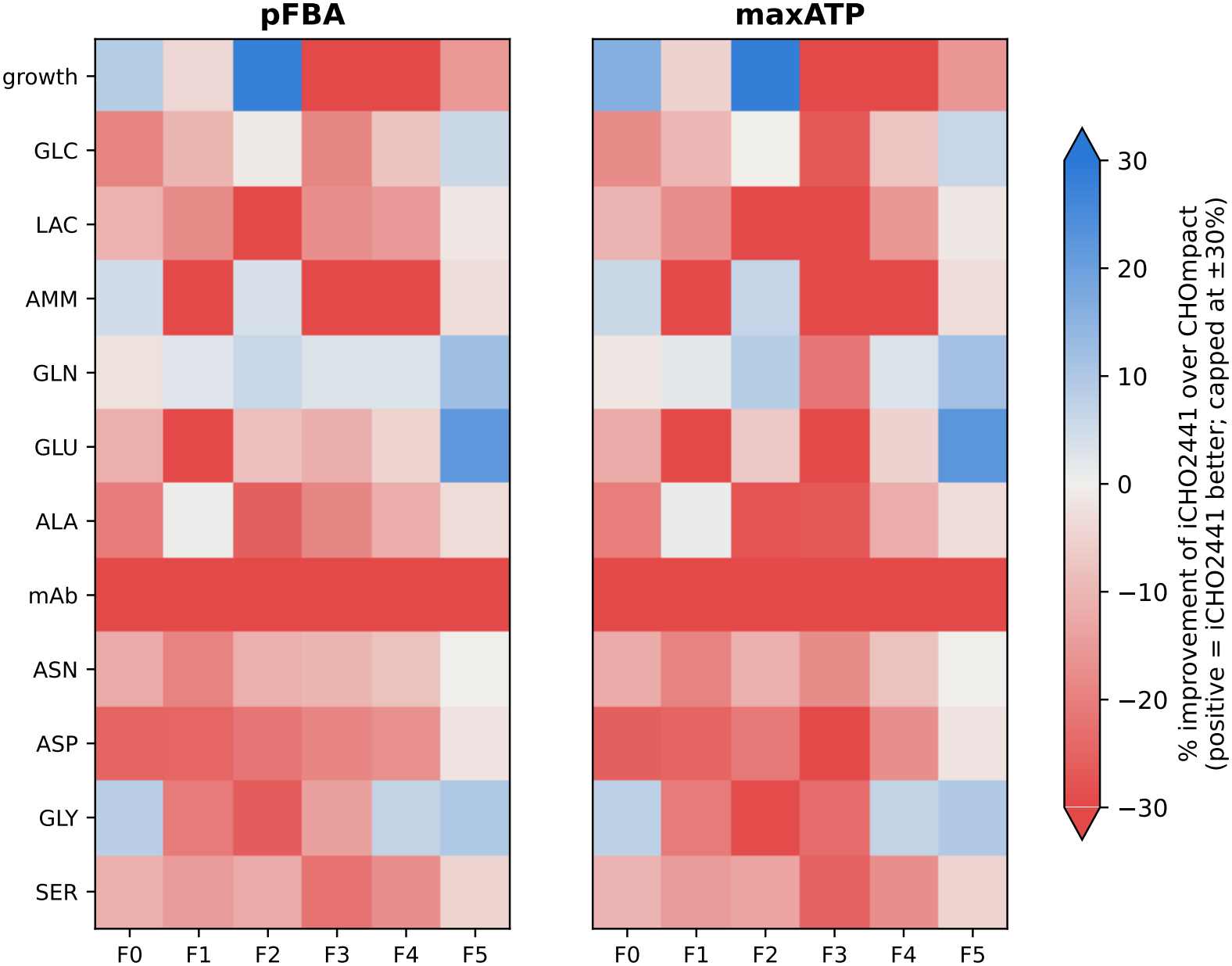
Resolution gain between model sizes for the two solution strategies (in exchange fluxterms), for each fold. A positive % of improvement (blue) signifies that that meta bolite is better described in iCHO2441.

CHOmpact proves advantageous over iCHO2441 in most cases, most clearly for mAb. Asparagine, aspartate and serine - already flagged in the previous section as less well predicted - also benefit most from CHOmpact. These results suggest that increasing model size, without a corresponding increase in measured intracellular fluxes or at least tighter bounds, offers little benefit to prediction accuracy. Both panels look similar, again indicating insensitivity to the choice of optimisation strategy.

iCHO2441 has 44× the number of reactions of CHOmpact, but once the model is built and the training setup is complete, it costs only ∼2.6× more compute per epoch (20.5 ± 0.4 vs 7.9 ± 0.3 ms) and ∼2.7× more memory (+46 vs +17 MB peak RSS), a markedly sub-linear scaling in both (Supplementary Table 5). The increase in computational cost is therefore not, on its own, a reason to prefer a reduced model, unless flux sampling is required. The case for CHOmpact rests instead on comparable prediction accuracy and more tractable flux interpretation, given the lack of measurement support for iCHO2441’s added reactions.

**Table 3:**
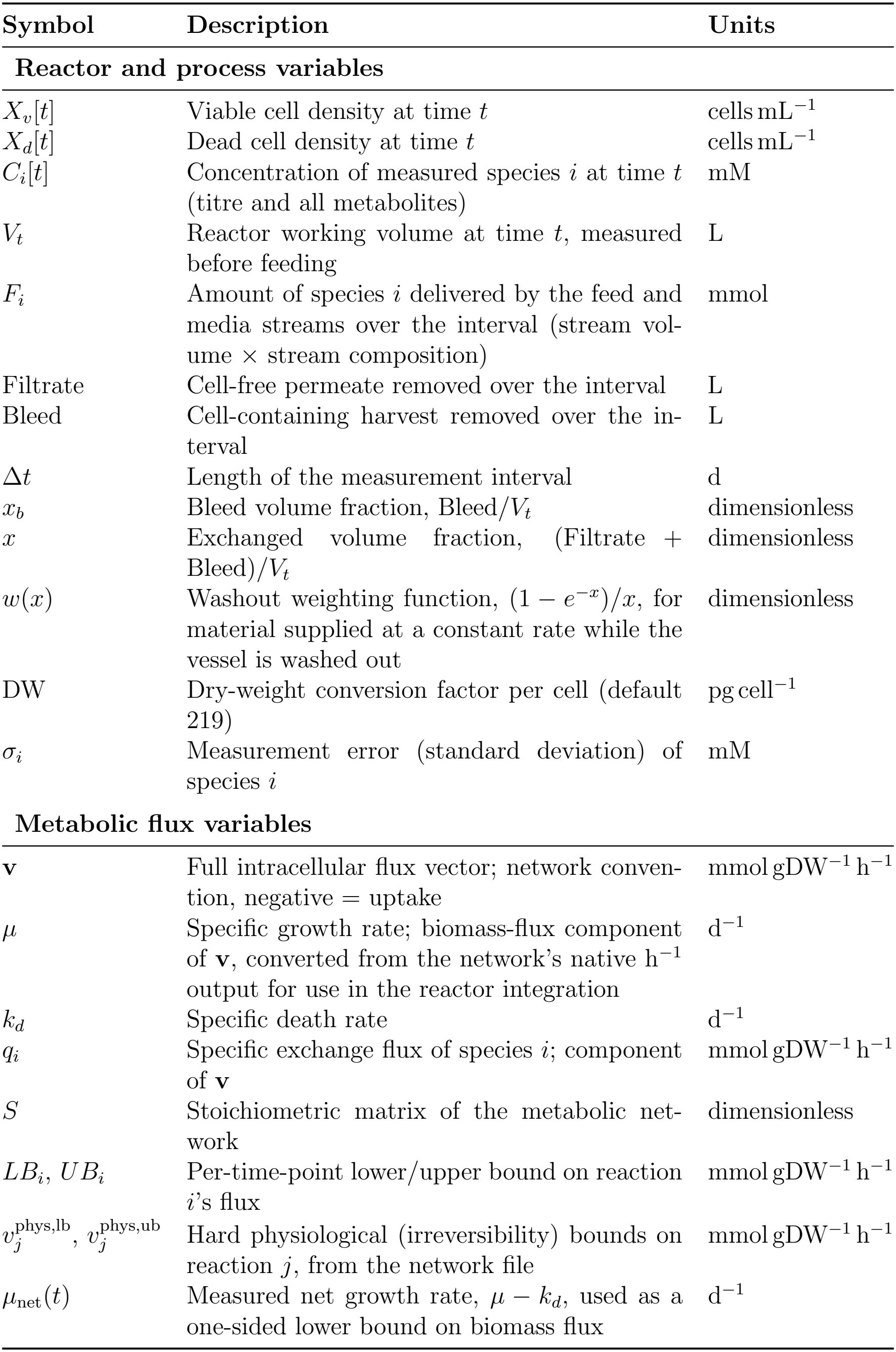
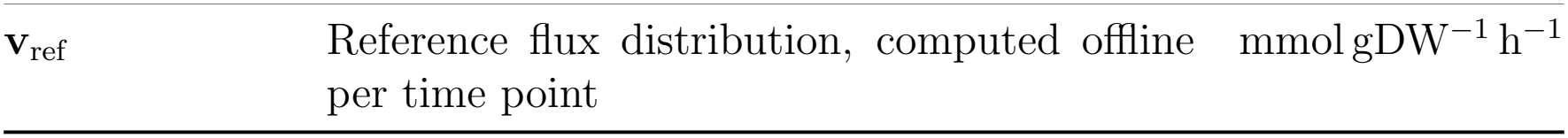
Nomenclature for the reactor and metabolic flux quantities.

| Symbol | Description | Units |
|---|---|---|
| **Reactor and process variables** |  |  |
| *X*<sub>*v*</sub>[*t*] | Viable cell density at time *t* | cells mL<sup>−1</sup> |
| *X*<sub>*d*</sub>[*t*] | Dead cell density at time *t* | cells mL<sup>−1</sup> |
| *C*<sub>*i*</sub>[*t*] | Concentration of measured species *i* at time *t* (titre and all metabolites) | mM |
| *V*<sub>*t*</sub> | Reactor working volume at time *t*, measured before feeding | L |
| *F*<sub>*i*</sub> | Amount of species *i* delivered by the feed and media streams over the interval (stream vol-ume × stream composition) | mmol |
| Filtrate | Cell-free permeate removed over the interval | L |
| Bleed | Cell-containing harvest removed over the in-terval | L |
| Δ*t* | Length of the measurement interval | d |
| *x*<sub>*b*</sub> | Bleed volume fraction, Bleed/*V*<sub>*t*</sub> | dimensionless |
| *x* | Exchanged volume fraction, (Filtrate + Bleed)/*V*<sub>*t*</sub> | dimensionless |
| *w*(*x*) | Washout weighting function, (1 − *e*<sup>−*x*</sup>)/*x*, for material supplied at a constant rate while the vessel is washed out | dimensionless |
| DW | Dry-weight conversion factor per cell (default 219) | pg cell<sup>−1</sup> |
| *σ*<sub>*i*</sub> | Measurement error (standard deviation) of species *i* | mM |
| **Metabolic flux variables** |  |  |
| **v** | Full intracellular flux vector; network conven-tion, negative = uptake | mmol gDW<sup>−1</sup> h<sup>−1</sup> |
| *μ* | Specific growth rate; biomass-flux component of **v**, converted from the network's native h<sup>−1</sup> output for use in the reactor integration | d<sup>−1</sup> |
| *k*<sub>*d*</sub> | Specific death rate | d<sup>−1</sup> |
| *q*<sub>*i*</sub> | Specific exchange flux of species *i*; component of **v** | mmol gDW<sup>−1</sup> h<sup>−1</sup> |
| *S* | Stoichiometric matrix of the metabolic net-work | dimensionless |
| *LB*<sub>*i*</sub>, *UB*<sub>*i*</sub> | Per-time-point lower/upper bound on reaction *i*'s flux | mmol gDW<sup>−1</sup> h<sup>−1</sup> |
| *v*<sub>*j*</sub><sup>phys,lb</sup>, *v*<sub>*j*</sub><sup>phys,ub</sup> | Hard physiological (irreversibility) bounds on reaction *j*, from the network file | mmol gDW<sup>−1</sup> h<sup>−1</sup> |
| *μ*<sub>net</sub>(*t*) | Measured net growth rate, *μ* − *k*<sub>*d*</sub>, used as a one-sided lower bound on biomass flux | d<sup>−1</sup> |
| **v**<sub>ref</sub> | Reference flux distribution, computed offline per time point | mmol gDW<sup>−1</sup> h<sup>−1</sup> |

**Table 4:**
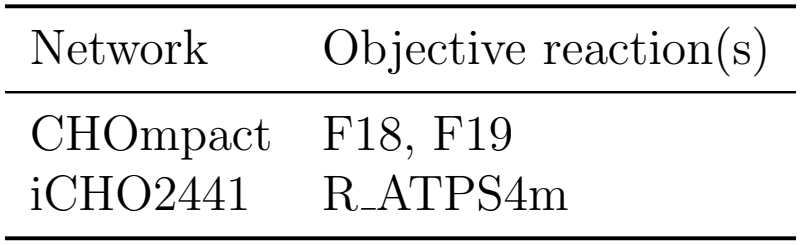
Reactions used as the max ATP objective for each metabolic network.

| Network | Objective reaction(s) |
|---|---|
| CHOmpact | F18, F19 |
| iCHO2441 | R_ATPS4m |

**Table 5:** Computational performance across network sizes. Mean ± std wall time per training epoch (epochs 2-20 of 20; five fed-batch experiments, batch size 4) and peak resident-set increment over the loaded process, single BLAS thread, Apple M5.

| Network | Reactions | Mean epoch time | Peak RSS |
|---|---|---|---|
| CHOmpact | 144 | 7.9 ± 0.3 ms | +17 MB |
| iCHO2441 | 6337 | 20.5 ± 0.4 ms | +46 MB |

#### Central-carbon flux predictions converge across metabolic optimisation strategies

As discussed in the previous Section, exchange flux predictions are relatively insensitive to solver choice, for both metabolic network sizes. The data-driven loss terms and stoichiometric feasibility constraints together anchor the exchange fluxes regardless of the optimisation objective.

To assess which solution strategy produces biologically coherent intracellular flux predictions, we examined predicted fluxes for a curated set of central-carbon reactions spanning upper glycolysis, the pyruvate node, the TCA cycle, anaplerosis, pentose phosphate pathway (PPP) and oxidative phosphorylation (OXPHOS), in both exponential phase (days 0-4) and stationary phase (days 9+) of the fed-batch dataset (Figure 8 for CHOm-pact and Supplementary Figure 18 for iCHO2441). The list of reactions included in each subsystem can be found in Supplementary Table 6 for CHOmpact and 7 for iCHO2441.

**Table 6:**
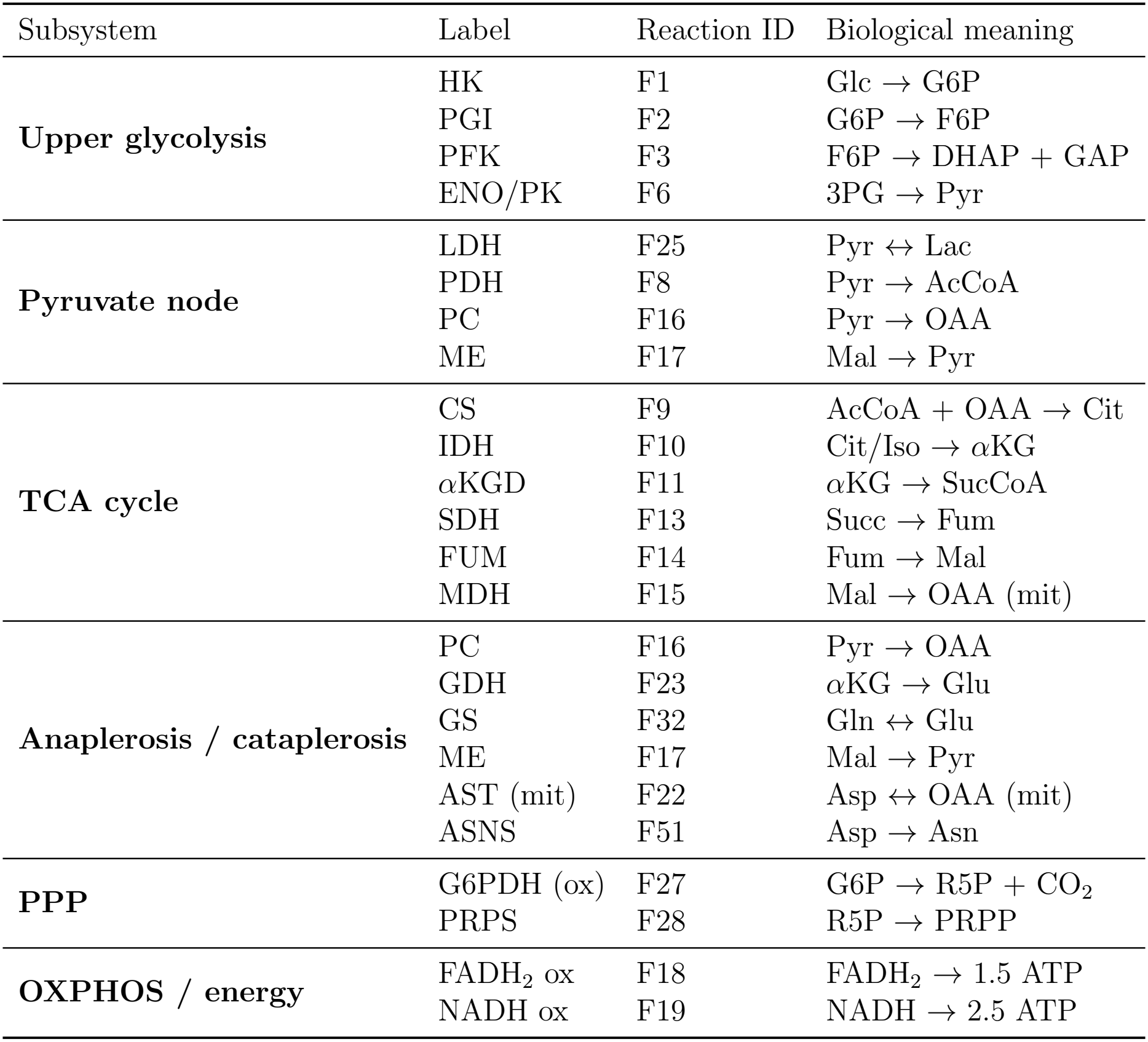
CHOmpact reaction map (F-numbers).

**Figure 8:**
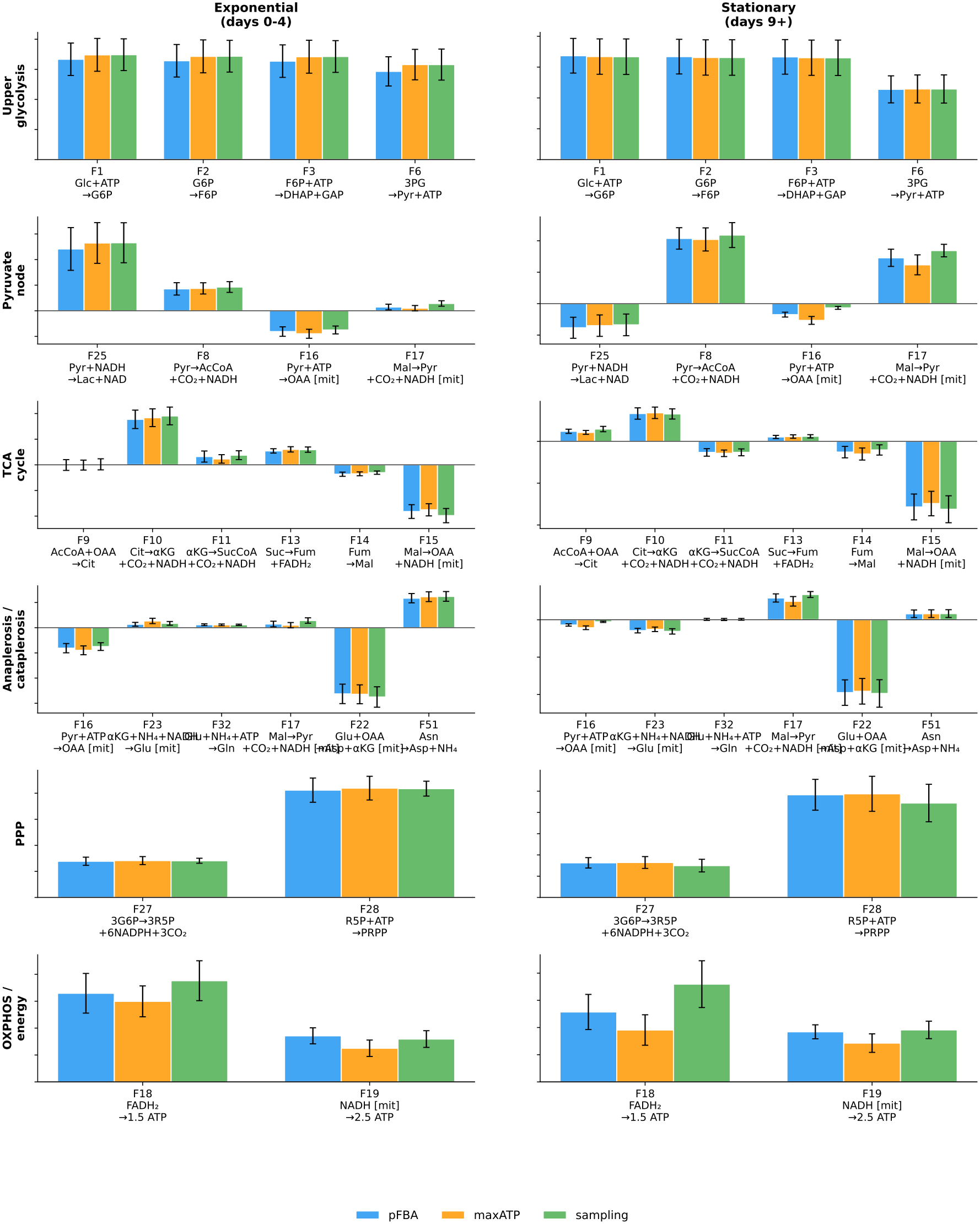
Intracellular fluxes for the three metabolic solution strategies (pFBA, max ATP and flux sampling) (model = CHOmpact).

In CHOmpact at exponential phase, the three solvers agree closely not only in upper glycolysis but throughout the pyruvate node, TCA cycle and anaplerosis. Across the curated reactions in these subsystems, pFBA, max ATP and flux sampling differ by less than their cross-fold standard errors in almost every case (Figure 8; values not shown). All three strategies also agree on the direction of F16 (Pyr + ATP → OAA [mit]) and F17 (Mal → Pyr + CO_2_ + NADH [mit]). Within these subsystems, the largest differences in magnitude between strategies are in flux F17, which is higher under sampling than under pFBA and max ATP, and in flux F23 (*α*KG + NH_4_ + NADH Glu → v[mit]), which is higher under max ATP than under pFBA, though its contribution to the network is small. In OXPHOS, max ATP consistently yields the lowest fluxes of the three strategies, although the differences are within cross-fold error. F18 (FADH_2_ → 1.5 ATP) and F19 (NADH[mit] → 2.5 ATP) are lowest under max ATP in both phases, and combined OXPHOS ATP output under max ATP is roughly 15-20% lower than under pFBA and sampling in exponential phase, widening to roughly 25-35% lower in stationary phase. This difference is systematic across cross-validation folds but small, and its ordering is opposite to that of the reference fluxes. The max ATP reference shows the highest OX-PHOS ATP output of the three strategies, whereas the predicted OXPHOS fluxes remain well below all three references, indicating that the reference term only weakly constrains these reactions. The ordering between strategies is therefore not interpreted metabolically. In stationary phase (second column of Figure 8), OXPHOS ATP output falls by roughly five-to six-fold relative to exponential phase under all three strategies, tracking the reduced glycolytic and TCA flux in this phase.

To assess whether the predicted fluxes are plausible, we quantified OXPHOS ATP yield per unit glucose consumed (computed as F18 × 1.5 + F19 × 2.5 from the lumped stoichiometry, divided by the glucose uptake flux F1). In CHOmpact at exponential phase this gives 1.38 (pFBA), 1.09 (max ATP) and 1.38 (sampling) ATP per glucose; in stationary phase, 0.50, 0.39 and 0.61 respectively. All three strategies sit well below the theoretical maximum of ∼30 ATP/glucose for full aerobic oxidation [31], consistent with the extensive aerobic glycolysis (lactate fermentation) reported experimentally in CHO culture, with no single strategy relying disproportionately on non-glucose carbon.

In iCHO2441 (Supplementary Figure 18), however, ATP synthase (ATPS4m) comes up as consistently negative under both solvers and phases, which is the wrong direction for a respiring, growing cell.

Taken together, these comparisons indicate that solution strategy choice is a minor source of uncertainty in intracellular flux predictions. The overall agreement between strategies partly reflects how weakly the unmeasured fluxes are constrained, by both the data and the reference term in the loss function. When differences are observed between the three strategies, they are small and do not follow the ordering of the reference fluxes, so they do not reflect a substantial difference in how each strategy routes carbon or energy. In CHOmpact, all three strategies also converge on a biologically plausible ATP yield. The more consequential factor is therefore network resolution relative to the data available to constrain it. CHOmpact’s smaller, more lumped representation keeps every reaction examined here, including OXPHOS, within a sensible range under any strategy. On the other hand, iCHO2441’s more granular representation resolves several of the same processes into unmeasured reactions and these are free to take implausible values independent of solver. Solver comparison and network resolution are therefore separate questions, and it is the second that currently limits what can be validated in iCHO2441.

#### Metabolic loss structure transfers from fed-batch to continuous

TL from fed-batch to continuous operation was evaluated under 3-fold leave-one-out cross-validation, with each fold training on six fed-batch experiments and two continuous experiments and predicting the held-out continuous experiment. We compared the different TL strategies, varying which parameter subsets of the trained fed-batch model are adapted during the continuous fine-tuning stage, as defined in Section Step 2: Fine-Tune Continuous Model.

**Figure 9:**
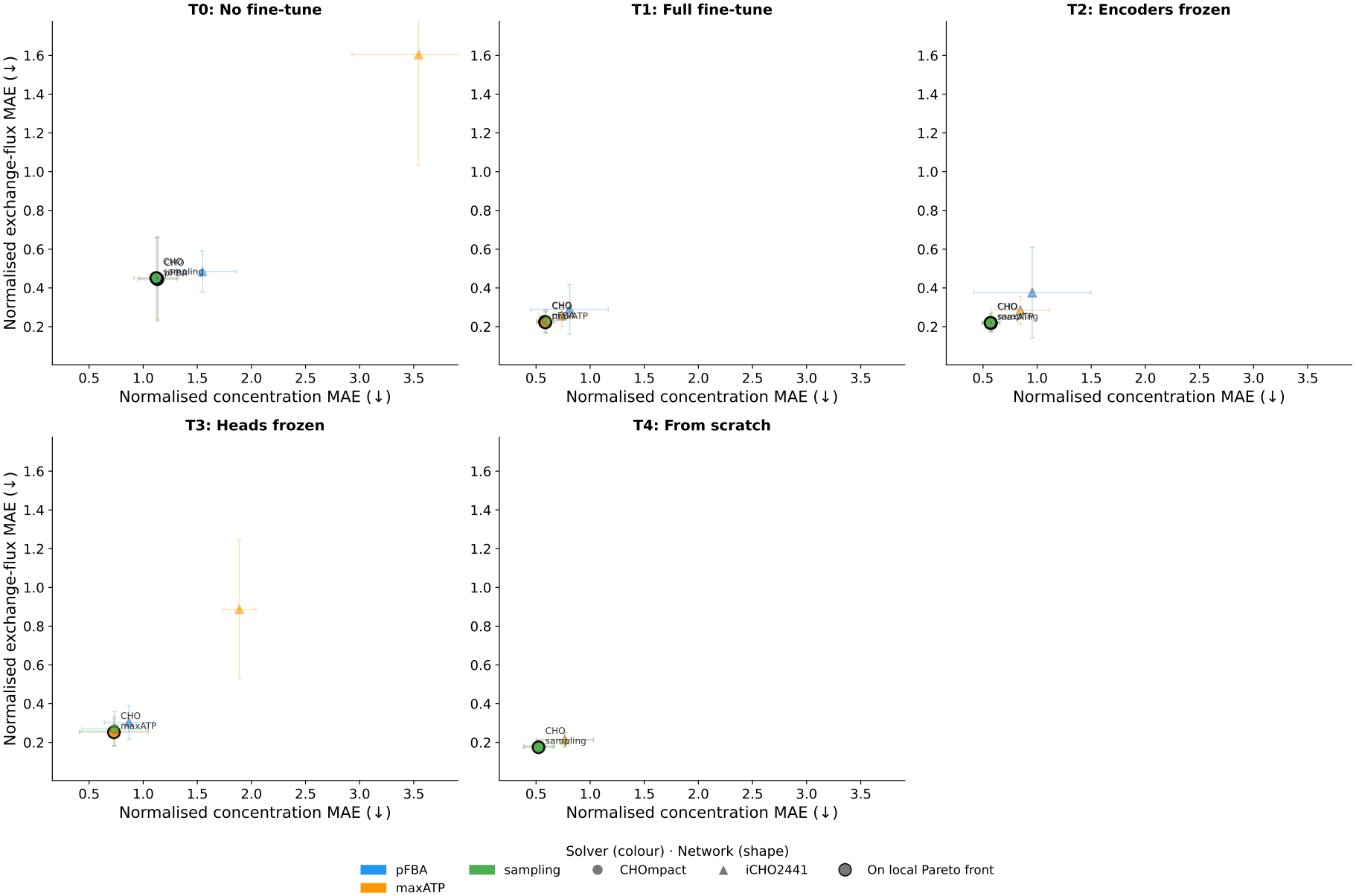
Pareto-front between concentration and flux prediction for different TL strategies.

Most combinations of optimisation strategy and network size cluster together at around a normalised MAE between 0.5 and 1.5 pertaining to concentration and between 0.2 and regarding exchange fluxes. An exception is the max ATP strategy with the iCHO2441 model size which is the least accurate in TL strategies T0 and T3.

No fine-tuning (strategy T0) shows the worst overall accuracy and the largest spread, while all remaining strategies achieve lower concentration and exchange flux normalised MAE. This constitutes the largest regime mismatch of any strategy, so it is expected to perform the worst.

Freezing the encoders (T2) and freezing the heads (T3) partition the network parameters, so together they isolate whether it is the input representation or the flux mapping that needs to adapt. T2 (encoders frozen, heads fine-tuned) performs close to T1, while T3 (heads frozen, encoders fine-tuned) retains the max ATP/iCHO2441 outlier seen in T0. This holds even though T2 keeps the encoder weights for the continuous-only streams (media, bleed and filtrate) at their initial values, whereas T3 learns them. Since the head is what maps the shared latent representation onto the flux vector, this suggests that adapting the head, rather than the encoder, is what corrects the mismatch for this configuration.

The structure of the metabolic losses in the metabolism-informed model is invariant to the mode of operation; on the other hand, the reactor kinetics (uptake/secretion rates, dilution) are not. In continuous mode, these differ substantially from fed-batch, in particular dilution. TL T1 retrains every tunable parameter. This is why the strategy performs as well as strategy T4 (training from scratch). The latter performs well most likely due to the small size and low variability of the continuous dataset rather than generalisation, and points to overfitting. The overall good performance of T1 additionally suggests that NEXT-FBA bounds transfer well between regimes.

Error bars are relatively wide across almost every panel, so within a given strategy, the differences among metabolic solution strategies and networks are mostly within error. This spread stems from the limited amount of continuous training data: with only three experiments, each fold trains on two, making the fit sensitive to which experiments the model sees.

Figure 10 shows the full fine-tuning (T1) model performance on the held-out experiments. Fold 2 appears as a systematic outlier, as it diverges from folds 0 and 1 for almost every metabolite, most notably overpredicting viable cell density. With only three continuous runs in total, one outlier is driving most of the disagreement.

**Figure 10:**
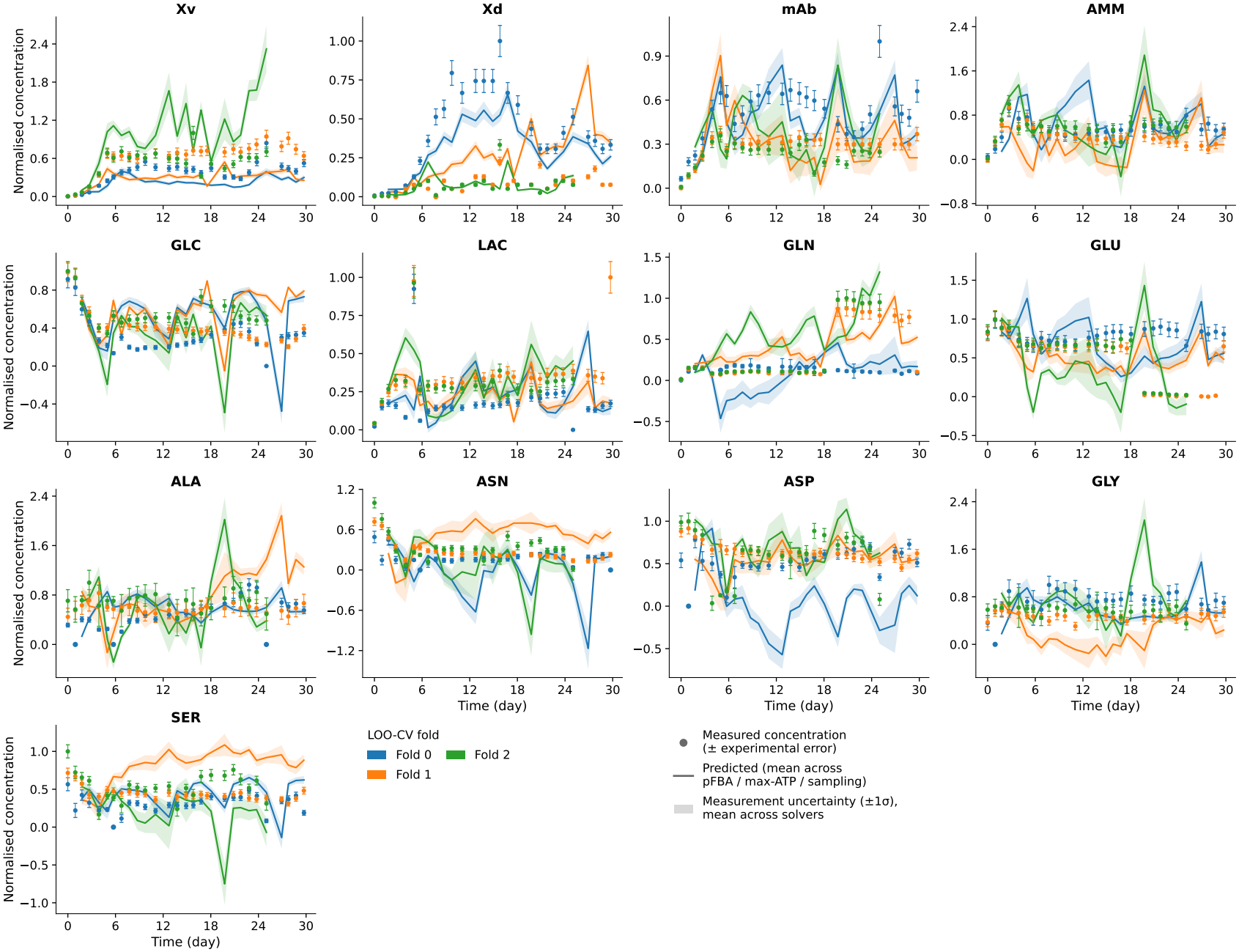
Concentration trajectory predictions (mean across solvers) for cross-validation experiments in continuous mode (model = CHOmpact, TL strategy = T1 Full fine-tune).

mAb, ammonia and glucose are tracked reasonably well across all three folds. As-paragine, aspartate, glutamate and to a lesser extent serine are predicted below zero. Asparagine and serine have been flagged in a previous Section for showing higher error in predicted fluxes, especially higher-magnitude fluxes, which is consistent with the difficulty of estimating growth under continuous culture. At steady state, viable cell density (VCD) is held constant while cells are continuously growing and dying, so only the net rate of change is observable from measured viable and dead cell density, and the true growth and death rates are not individually identifiable. As a result, the metabolism-informed model struggles to satisfy nutrient uptake and metabolite secretion under little apparent growth. An overpredicted flux that is propagated to the reactor mass balances, with the dilution effect that occurs in continuous mode, will amplify the error into the concentration prediction. The non-negativity penalty is not a hard constraint and it proves to be insufficient to deal with this case. The dilution term itself is a second source of error here: the harvest rate was back-calculated from reactor volume, media addition and bleed rates because measured and calculated volumes did not match (Section Experimental data), adding noise to the same dilution term implicated above. The uncertainty band is only propagating measurement error, so it does not capture this model-driven, flux-originating uncertainty either.

#### Model captures the lactate shift and predicts higher metabolic efficiency in continuous operation

The proposed metabolism-informed model has been shown to produce accurate metabolite concentrations and fluxes in fed-batch mode. The proposed model structure has also been used to characterise the methodological modelling trade-offs when transferring to continuous mode of operation in low data environments. Herein, we will show how this model formulation can be useful to generate hypotheses about how CHO cell metabolism differs between fed-batch and continuous.

Principal component analysis (PCA) on the measured exchange fluxes in fed-batch and continuous mode suggests that growth might be the dominant axis of variation between exchange fluxes and not mode of operation, as it is the main distinguishing factor between exchange fluxes. Continuous-mode fluxes cluster with fed-batch late-stage fluxes rather than with growing fed-batch fluxes (Figure 11, PC1 explains 56% of the variance of the data, while PC2 32%). However, when looking at the PCA loadings (Supplementary Figure 19), PC1 is largely driven by alanine flux (loading +0.88), with secondary contributions from asparagine ( −0.36), serine (− 0.20), and aspartate (− 0.19). The growth flux itself contributes little to PC1 as the separation of growing time points is carried by the amino acid fluxes that change with growth phase. PC2 (32%) is driven by a single high-glycine time point in fed-batch growth (day 1) and as such, it is excluded from interpretation. Fed-batch time points in exponential phase (0–4 d) separate from non-growing fed-batch and continuous time points along PC1. This is supported by earlier findings that report no significant differences in TCA cycle activity, glycolytic/oxidative split at a known branch point, or dedication to the PPP at the glucose-6-phosphate (G6P) branch point between fed-batch and continuous mode of operation [32]. Since continuous steady-state and fed-batch decline time points occupy the same low-PC1 region, we restrict the analysis to matched non-growth physiological states: fed-batch decline (*>*8 d) and continuous steady-state (5-15 d) (Figure 12).

**Figure 11:**
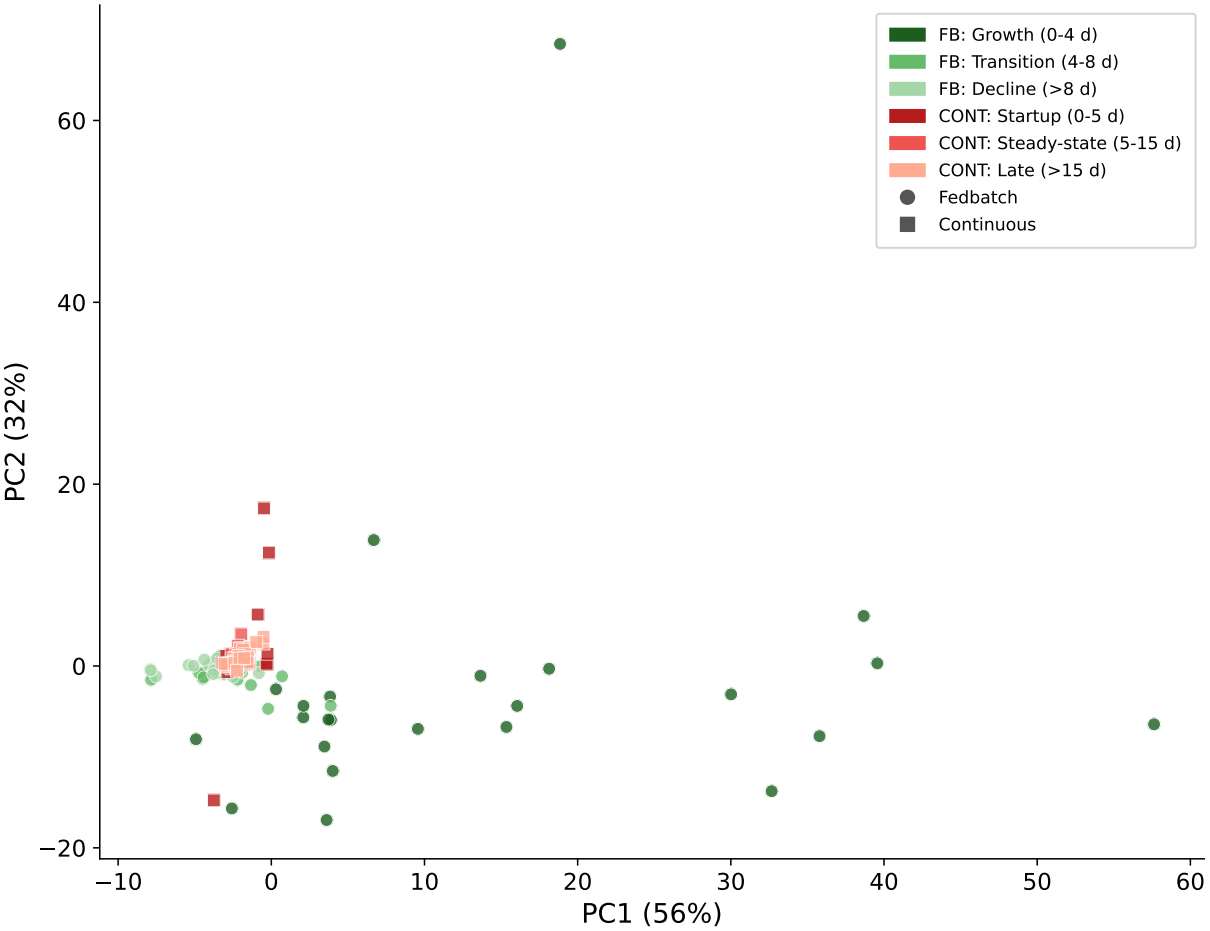
PCA on measured exchange fluxes. Different colours refer to different cell culture phases, circles refer to fed-batch and squares to continuous. Note: PCA was not run on raw flux data; instead they were scaled using scikit-learn’s RobustScaler (median-centred, interquartile-range scaled), therefore the loadings do not reflect magnitude of the fluxes.

**Figure 12:**
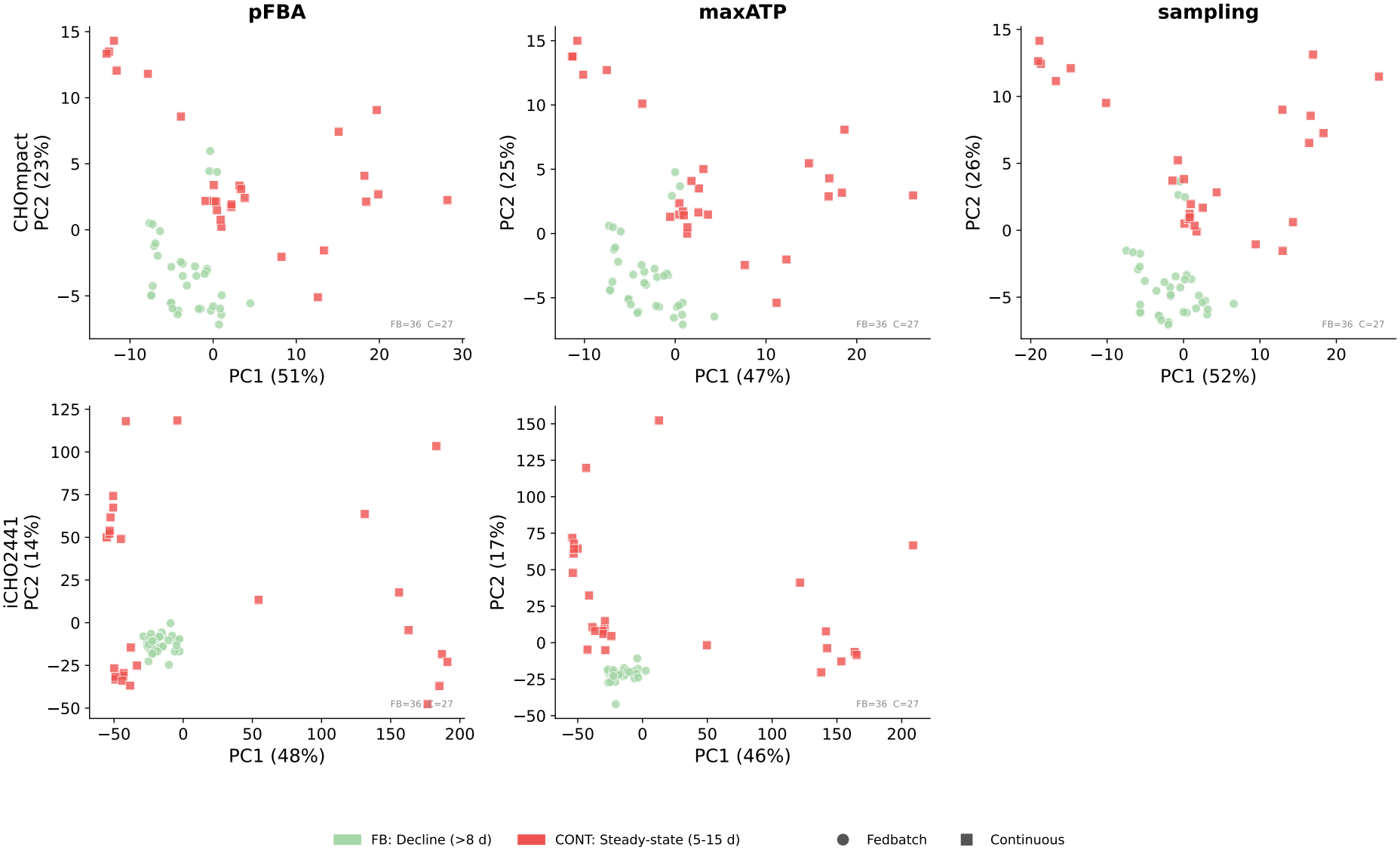
PCA per continuous model, controlled for growth. (model = CHOmpact and iCHO2441, TL strategy T1)

CHOmpact model (top row of Figure 12) across all three solvers presents visible separation between fed-batch decline (green) and continuous steady-state (red) along PC1. The strategy max ATP shows the cleanest separation, but pFBA and sampling also show some degree of separation with higher scatter. The separation between modes of operation is less visible in the iCHO2441 model (bottom row).

Given that different model configurations have been shown to vary with methodological choices (notably TL strategy), we investigate herein if there are regime-shift predictions which are robust to those choices, and, therefore can be made more general. Fifteen configurations were run corresponding to three solution strategies times five TL strategies, for the CHOmpact model. For each reaction in each configuration the difference between mean predicted flux in continuous and fed-batch modes of operation was computed in exponential and stationary phases (Figure 13). The reactions were sorted according to the most meaningful flux difference between fed-batch and continuous modes. Bars show the mean flux difference and error bars the standard deviation across the 15 configurations. Reactions are restricted to those with a robust difference. To be considered robust, the mean flux difference must exceed a data-driven magnitude threshold, the sign of the difference must agree in ≥90% of the 15 configurations, and the relative variability must be *<*0.5. Hatched bars with red reaction labels and an asterisk denote bound-violating reactions, for which fewer than 90% of pooled time points (across all configurations, in at least one of the two compared phases) satisfy *v* ≥ *LB* − 0.01; these are flagged rather than excluded because the model’s architecture hard-enforces only the mass-balance constraint (*S***v** = 0), while flux bounds are imposed solely as a soft training penalty.

**Figure 13:**
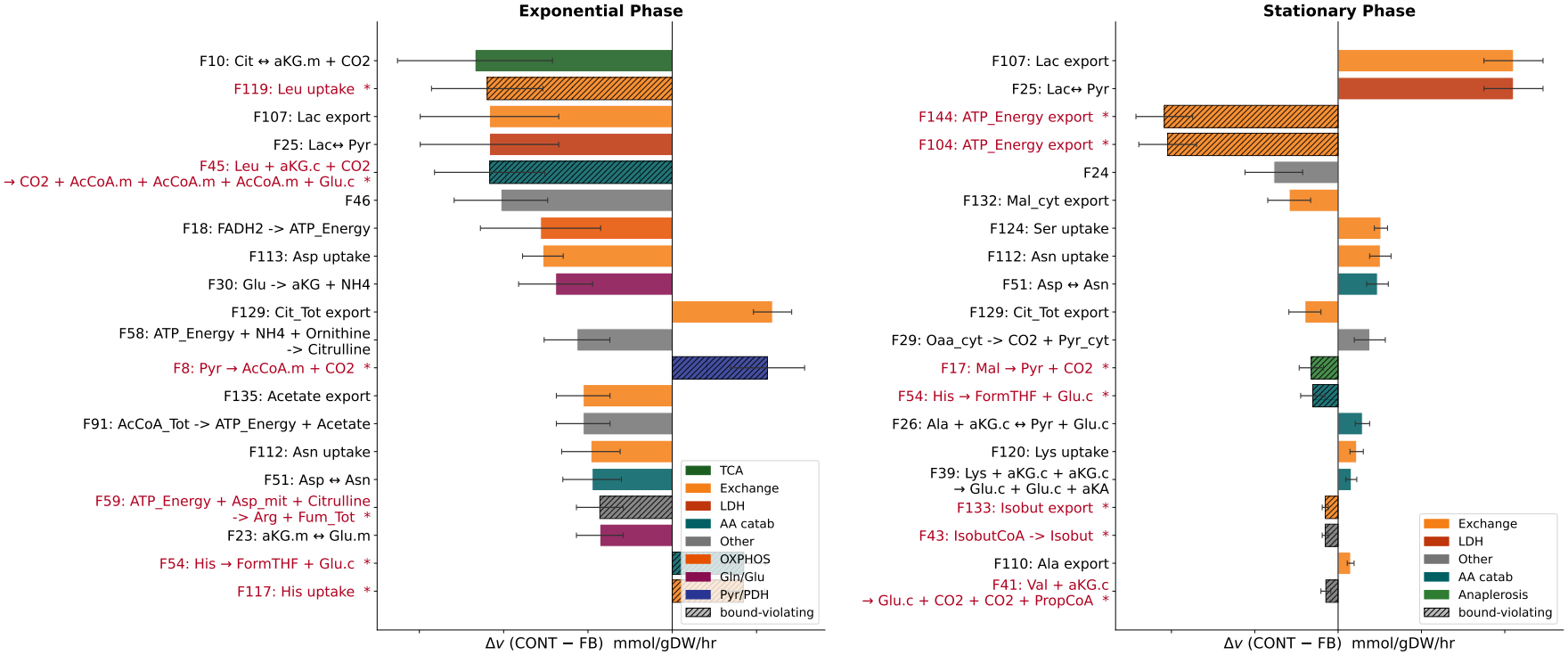
Predicted intracellular flux shifts between continuous and fed-batch culture in CHOmpact model. Bars show the mean change in predicted flux for the top 20 reactions ranked by largest difference in the exponential phase (left; fed-batch Growth ≤4 d vs. continuous Startup ≤5 d) and stationary phase (right; fed-batch Decline *>*8 d vs. continuous Steady-state 5–15 d). The flux difference was computed for each of 15 model configurations (3 metabolic solvers – pFBA, max ATP, flux sampling – × 5 transfer-learning strategies T0–T4; continuous configurations pool the three leave-one-out validation folds), using the direct, unfiltered model flux predictions.

**Figure 14:**
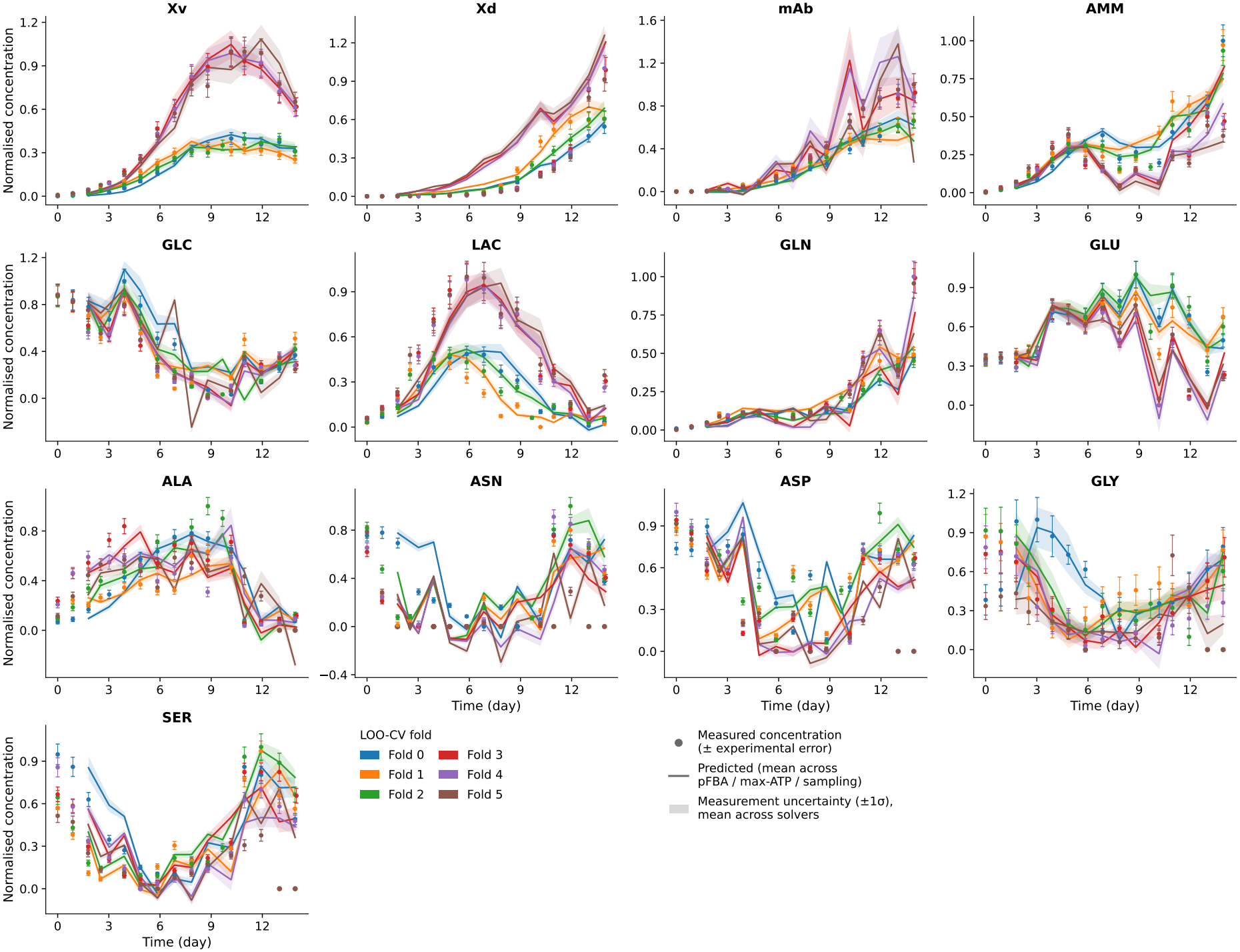
Concentration trajectory predictions for cross-validation experiments in fedbatch mode (model = iCHO2441). Each solid line is the mean prediction across pFBA, max ATP, and flux sampling.

**Figure 15:**
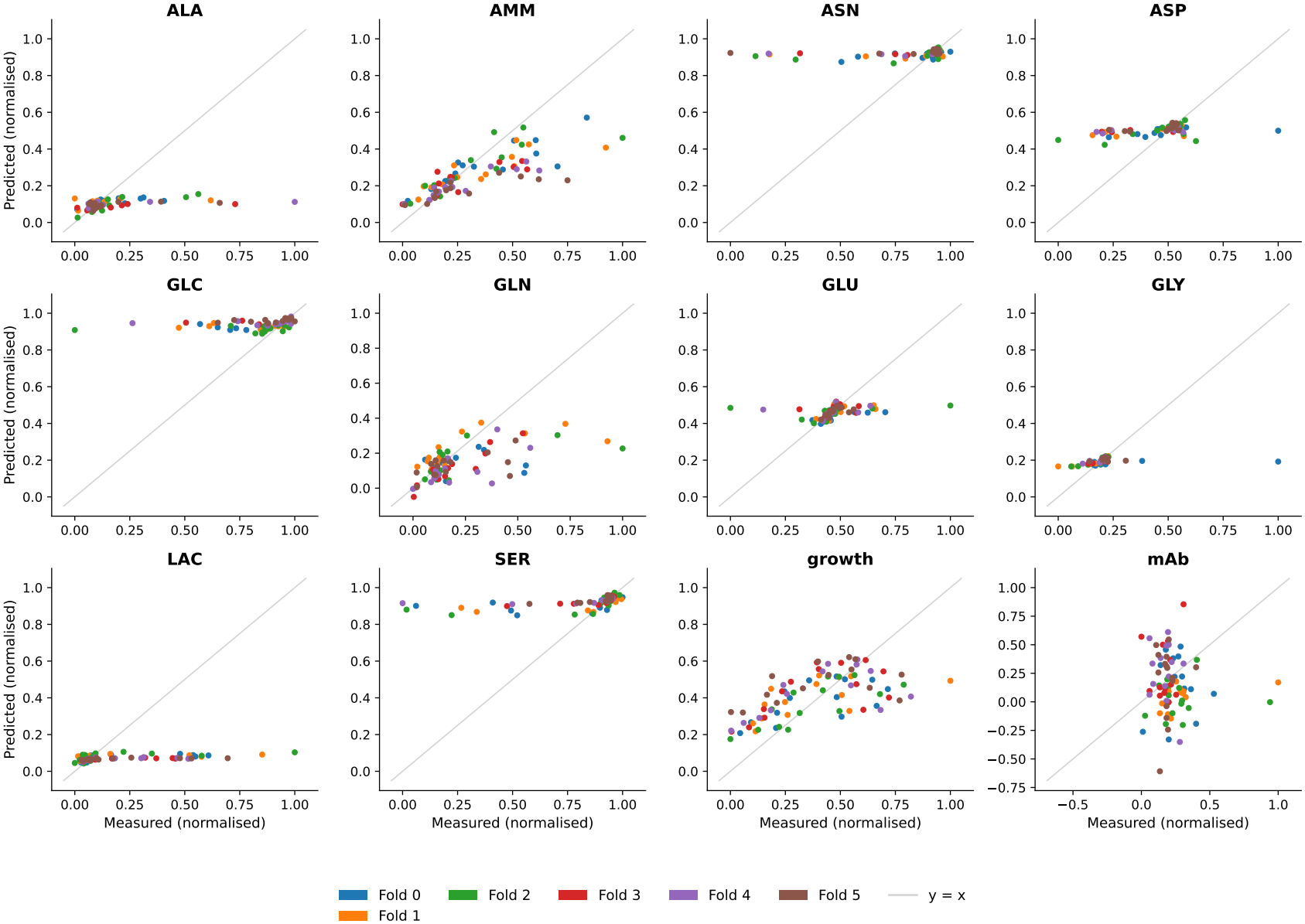
Exchange flux predictions for cross-validation experiments in fed-batch mode (model = iCHO2441).

**Figure 16:**
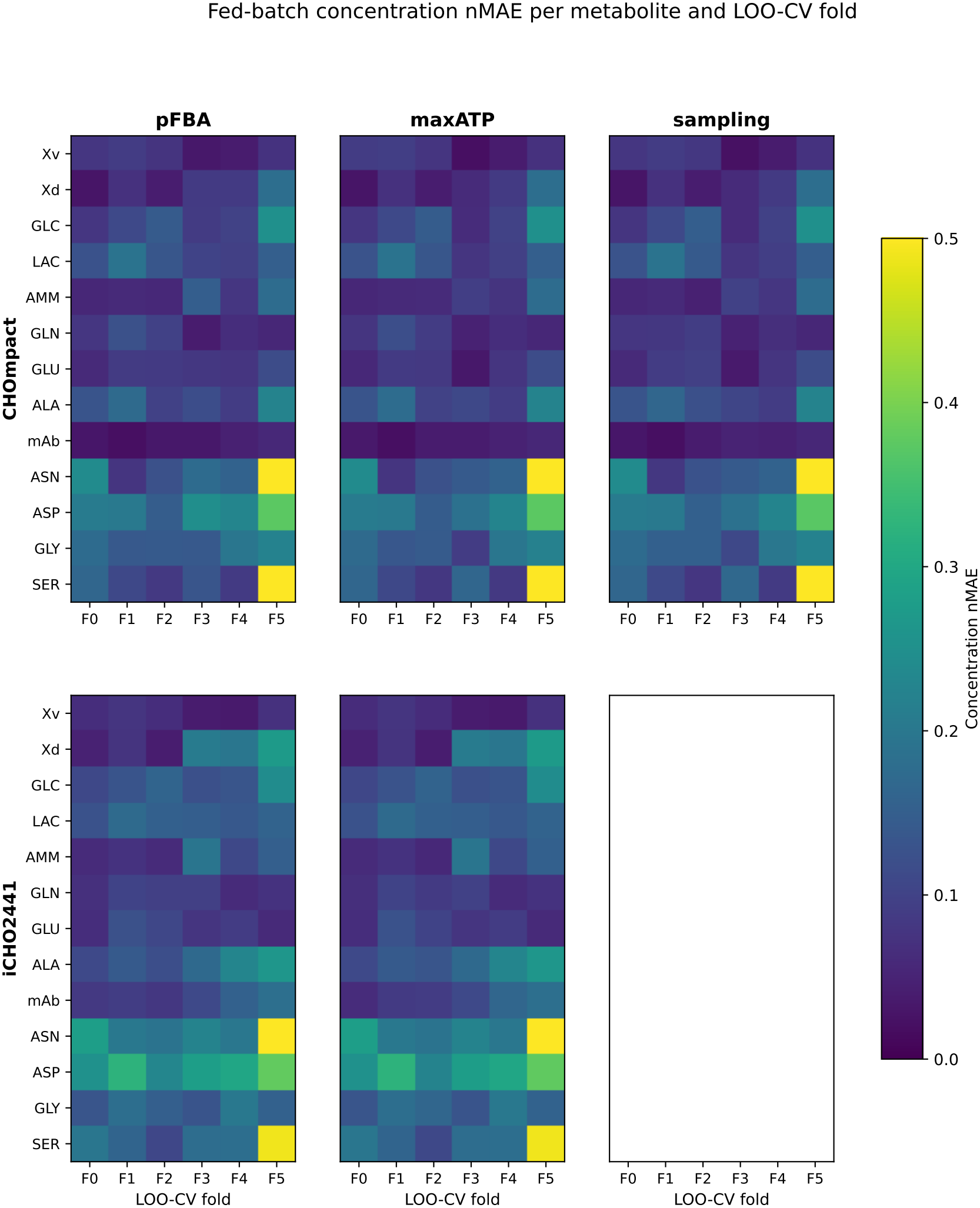
Normalised MAE of metabolite concentration, across all metabolic network model sizes and solution strategies.

**Figure 17:**
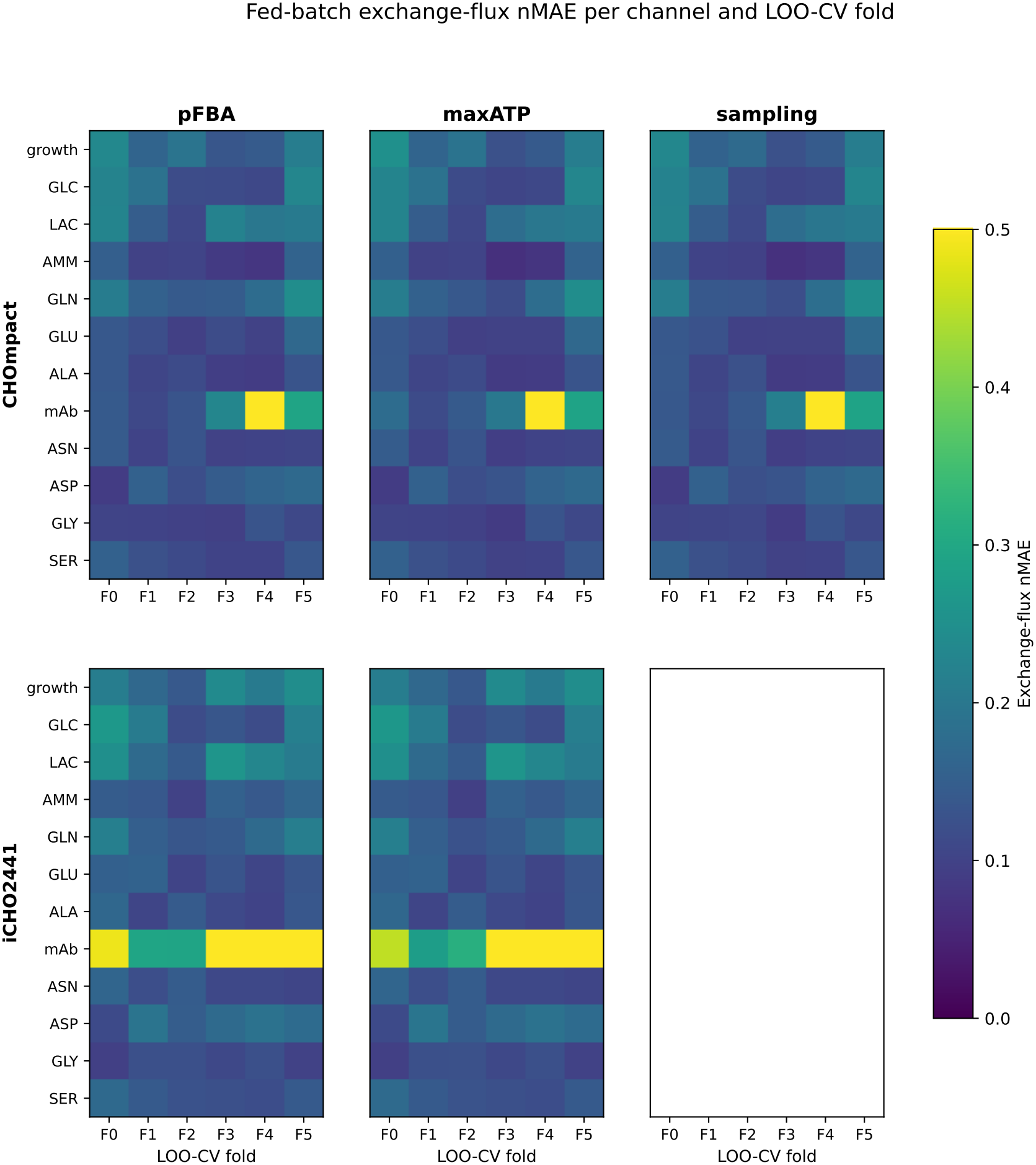
Normalised MAE of exchange flux prediction, across all metabolic network model sizes and solution strategies

**Figure 18:**
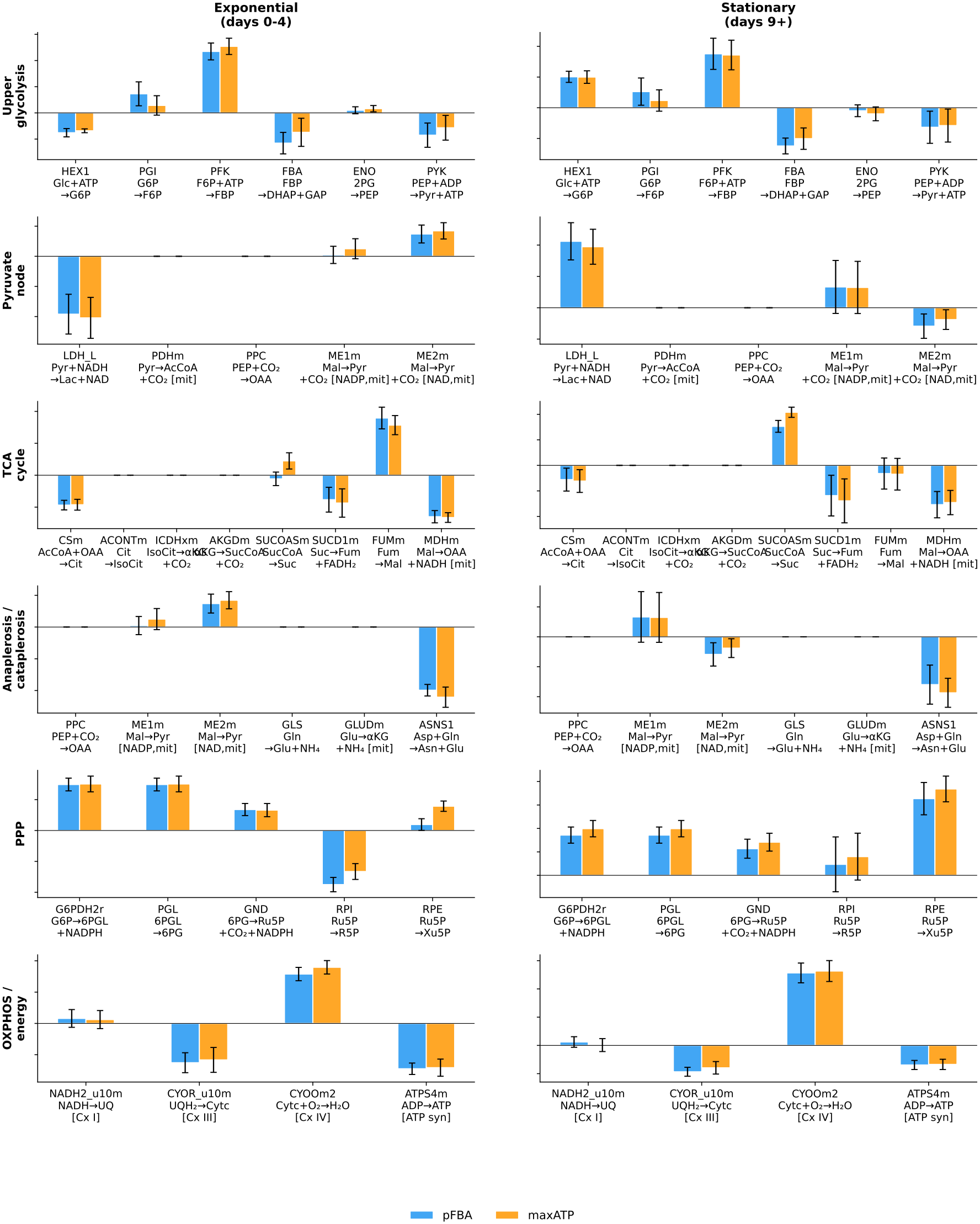
Intracellular fluxes for the metabolic solution strategies (pFBA and max ATP) (model = iCHO2441).

Figure 13 shows how the predicted intracellular fluxes differ between continuous and fed-batch operation in each phase. These shifts are not measured, but are instead a result of how the metabolic model accommodates the predicted exchange flux shifts, given the solution strategy and bounds. The reactions shown therefore represent intracellular adaptations that the model places consistently across solver and TL choices. Since the difference in flux is defined as continuous minus fed-batch, negative bars indicate reactions carrying more flux in fed-batch. In exponential phase most of the differences are negative, whereas in stationary phase the direction reverses.

The largest exponential-phase differences are roughly double the largest stationary-phase differences. This is partly because some of the biggest differences stem from measured exchange fluxes (F107, F113, F112). These are the reactions most directly driven by extracellular concentrations and dilution terms in the reactor mass balances. It is sensible that these are the top fluxes to diverge. On top of this, it has been discussed in previous Sections that with the current setup, it is not possible to calculate true growth rate. This means that while cells are growing and dying, the metabolic model struggles to reconcile an apparent flat growth rate with uptake and secretion of metabolites. As a result, reactions most connected to growth will be flagged with the biggest differences.

While both modes of operation show similar growth rates, in fed-batch, cells take up more glucose, asparagine, aspartate and serine and secrete more lactate, ammonia and alanine. This is why the core central carbon metabolism reactions such as F10, F25, F107, F18 and F30 are amongst the top differences between the two modes. Taken together, this suggests cells in continuous mode are more efficient for the same growth.

In stationary phase, the comparison is different because while in fed-batch cells are starting to decline, in continuous they are at steady state. This is why the major differences are reactions F25 and F107, which match measured reactions and are in line with the lactate shift expected at this stage of the culture.

Reactions highlighted in red violate their bounds, 6 in exponential phase and 7 in stationary phase (12 distinct reactions, as F54 appears in both). F8 (PDH), F54 (His catabolism), F117 (His uptake) in exponential and F144/F104 (ATP export), F17 (malic enzyme), F54, F41 (Val catabolism) in stationary violate their lower bounds in both modes of operation. On the other hand, F119 (Leu uptake), F45 (Leu catabolism), F59 (Arg synthesis) in exponential and F43/F133 (isobutyrate) in stationary only violate their bounds in continuous. These fluxes belong either to amino acid metabolism or to energy metabolism. These reactions are irreversible, meaning their lower bound is 0. Out of these 12 reactions, only four of them (F8, F17, F41 and F45) were constrained with an upper bound from NEXT-FBA. This means the remaining 8 were essentially unconstrained, as the bounds penalty is scaled by the window width, and a negative flux in a [0, 1000] window is barely penalised. Enforcing irreversibility as a hard constraint, or tightening the bounds with additional data, would be needed to address this violation.

The results suggest the model is routing resources in order to satisfy measured fluxes, through reactions that are not constrained by data. A negative flux through the ATP export reactions F104 and F144 would mean ATP is flowing into the cell from the out-side, which is not plausible. Negative fluxes through F8, F45 and F17 would mean they are consuming *CO*_2_, acetyl-CoA and NADH. Additionally, the model predicts cells are consuming aspartate and glycine and secreting glutamate. In order to abide by this, the model makes the cell secrete histidine, valine, threonine, tryptophan and leucine, using unmeasured fluxes to dispose of surplus nitrogen. This is a common occurrence in unconstrained networks, where the model uses these unconstrained fluxes to absorb any slack needed to satisfy the measured fluxes.

Literature reports higher branched-chain amino acid (BCAA) catabolism in continuous mode, in association with higher death rate [32], at steady state. BCAAs (Leu, Ile, Val) are not directly measured in our experiments, and are thus only predicted by the model. Still, none of these are flagged as a reliable source of differences at stationary phase, as the valine-related reactions that do appear (F41, F43, F133) all violate their bounds. Given what has been discussed about the reactions violating their bounds, measuring these amino acids would likely improve the model predictions substantially and help distinguish further the two modes of operation.

## Discussion

In this study, we describe a metabolism-informed model formulation that makes use of mass balances at reactor and cell level, coupled with a recurrent neural network architecture to represent a mammalian cell culture. The proposed model structure is capable of making predictions in continuous mode of operation, while being mainly trained with fed-batch data. To achieve this, the model architecture is agnostic to specific kinetic structures and modes of operation: the recurrent neural network predicts the fluxes directly rather than through predefined rate laws, so the structure holds regardless of operating mode. These fluxes are parameterised through the null space of the stoichiometric matrix, such that intracellular steady state holds by construction. The predicted fluxes are then passed through the reactor mass balance, which is integrated over each measurement interval without assuming steady state at the reactor level, as this is not observed in data. Metabolic knowledge is added through two soft penalties: one encourages the fluxes to stay within their bounds, with extracellular bounds from data and intracellular ones from NEXT-FBA [23], and the other pulls unconstrained reactions towards a reference flux distribution. This strategy was adopted for two practical reasons: it reduces the number of learned parameters, since only the null-space coordinates are predicted rather than the full flux vector, and it avoids embedding a metabolic solver in the forward pass, which could break backpropagation, and hence learning.

In Part I of the results, the weights contributing to the loss function were tuned with hyperparameter optimisation and benchmarked against a fully data-driven model architecture. In fed-batch mode, where more data is available, the metabolism-informed architecture led to comparable performance in both concentration and exchange flux prediction. When transferring to less abundant continuous data, the metabolism-informed model performed worse. Our main hypothesis is that the similarity across datasets benefitted a data-driven architecture, prone to overfitting. If slightly different data were to become available the data-driven architecture would likely underperform.

In Part II of the results, the contributions to the loss function were calibrated to the same magnitude as the data term. Two metabolic model structures and three metabolic solution strategies were benchmarked. Genome-scale model network size is shown to expand the feasible flux space available to the solver. This effect does not carry over to the concentration predictions at the reactor level, as exchange fluxes are tightly constrained by measured data. Although the additional computational cost of using a genome-scale model did not prove to be prohibitive (with the exception of flux sampling), the case for a reduced network rests on increased interpretability, given the lack of intracellular flux data. If transcriptomic or proteomic data are available, constructing a custom model would likely prove advantageous in terms of increasing the certainty of active pathways. The architecture of the metabolism-informed model makes it so that the solution strategy of the metabolic model matters little for intracellular interpretation even if less anchored by data. All three metabolic optimisation strategies yield similar flux values in major pathways. This makes the model architecture advantageous in the cases where there is interest in using a metabolic model but no sensible objective function is available.

This is because this model structure is able to combine the learned representations of a data-driven model with the interpretability of a metabolic model.

Using the model trained on fed-batch data and fine-tuning it with continuous data (TL strategy T1) demonstrated the highest agreement with data. Our hypothesis is that the metabolism-informed model, trained on six fed-batch experiments, has learned some metabolic relationships that are generalisable. Performing transfer learning with three continuous datasets allows for learning representations that are different (such as different feeding regimes). This is also why training from scratch (TL strategy T4) is equally satisfactory. However, similarly to the data-driven architecture, training from scratch on three continuous datasets is likely to lead to overfitting.

Still, the model performs worse in continuous than it does in fed-batch. Several factors could have contributed to this degradation besides lack of data. The first is that the growth and death rates are non-identifiable in our current continuous culture setup, so the model does not relate growth with uptake or secretion of any metabolites. Second, the measured and calculated volumes did not match, and this mismatch cannot be attributed to a single stream. We treated the harvest rate as the unreliable term and back-calculated it, but the bleed may also be inaccurate. Since cells are removed through the bleed only, this would add error to the viable cell density balance, and through the back-calculated harvest, to the metabolite balances as well. Lastly, the change of inputs to the model (in fed-batch there is no bleed and harvest rate for example) meant that the continuous-only inputs were never seen during fed-batch training, which could degrade the mapping between the two modes. One possible alternative to address this last point would be to use a different model architecture, such as autoencoders [33], as the common information would be compressed into a latent space and the different information would be the distinguishing features. However, it requires a substantial amount of data, which was deemed infeasible for this current application.

The performance of the transfer learning exercise suggests that a well-trained model on fed-batch data could aid in experimental design prioritisation for continuous culture characterisation. Alternatively, to achieve a satisfactory model making predictions in continuous mode, the growth and death rates need to be decoupled. An option could be to measure other markers, such as lactate dehydrogenase (LDH) or *O*_2_, which could serve as proxies for death rate. For this to be useful, these markers need to be measured in both regimes, otherwise it would further contribute to the misalignment of inputs in both modes.

PCA on measured exchange fluxes shows continuous steady-state time points cluster with late fed-batch, not with growing fed-batch, suggesting that growth rate rather than mode of operation is the primary axis of metabolic variation. We proceeded to analyse the mean predicted flux difference between continuous and fed-batch, ranking reactions by the magnitude of this difference across all 15 configurations (3 solvers × 5 TL strategies), separately for exponential and stationary phases. Several of the reactions in both phases originate from measured exchange fluxes and are therefore directly informed by data. The largest differences between fed-batch and continuous modes were observed in central carbon metabolism. In fed-batch, reactions carried substantially more flux during exponential phase, whereas in stationary phase, the opposite was observed. The intracellular fluxes reflected the metabolic steady-state characteristic of continuous mode of operation.

The main limitation of this study is data availability. While some mitigation measures were employed, such as dropout and the mechanistic priors, the extent to which they help the model make predictions in completely different data regimes remains untested in this study, as the total number of experiments is low and the diversity between them limited. Future work should prioritise increasing both quantity and diversity of training data through inclusion of additional cell lines and process conditions.

## Author Contributions

**Mariana Monteiro (MM)**: Conceptualisation, Methodology, Formal analysis, Investigation, Visualisation, Writing - Original Draft, Writing - Review & Editing. **Stacy Be-diako (SB)**: Investigation. **Ricardo Tecson (RT)**: Investigation. **Gregory Nierode (GN)**: Supervision, Writing - Review & Editing. **Pavlos Kotidis (PK)**: Supervision, Writing - Review & Editing. **Cleo Kontoravdi (CK)**: Conceptualisation, Methodology, Supervision, Writing - Review & Editing, Funding acquisition, Project administration.

## Acknowledgements

MM thanks the UK Biotechnology and Biological Sciences Research Council (BBSRC) and GSK for her studentship.

## Competing interests

The authors acknowledge the funding of this study by GSK through the BBSRC award.

## Data Availability

The data used in this study cannot be shared due to confidentiality reasons.

## Code Availability

The code used for this work will be available upon publication at https://github.com/marianaipmonteiro/MetabolismInformedModel.

## Glossary

Ala: alanine. 4
Amm: ammonia. 4
Asn: asparagine. 4
Asp: aspartate. 4
CHO: Chinese hamster ovary. 1, 3, 4, 17, 22
Glc: glucose. 4, 5
Gln: glutamine. 4
Glu: glutamate. 4
Gly: glycine. 4
GMP: Good Manufacturing Practice. 3
Lac: lactate. 4
LDH: lactate dehydrogenase. 27
LSTM: long short-term memory. 6, 8–11
mAb: monoclonal antibody. 1, 3, 4, 13, 16, 21
MAE: mean absolute error. 12, 13, 16, 19, 37, 40, 41
PCA: principal component analysis. 22, 23, 27, 46
Ser: serine. 4
TL: transfer learning. 9, 10, 19, 20, 22–24, 26, 27
VCD: viable cell density. 21

## Nomenclature

## Detailed Model Structure

### Metabolic Optimisation Strategy

For each valid time point *t*, all three metabolic optimisation strategies (parsimonious FBA, max ATP, and flux sampling) operate on the same feasible set. Mass balance is imposed exactly,

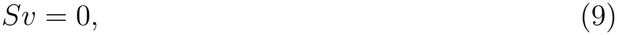

and fluxes are bounded by a relaxable, time-dependent window together with a hard physiological box:

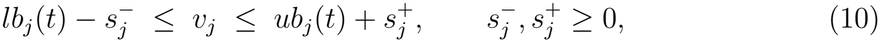

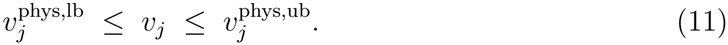

The slack variables 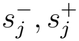 in Eq. (10) allow the window itself to be violated at a cost, whereas the bounds in Eq. (11) are never relaxed and enforce reaction irreversibility. The windows [*lb*_*j*_(*t*), *ub*_*j*_(*t*)] are the measured exchange windows for the extracellular fluxes, the intracellular bounds derived from NEXT-FBA [23] on mapped intracellular reactions, and the physiological box everywhere else. The biomass reaction receives a one-sided bound, *lb*(*t*) = max(0, *µ*_net_(*t*)).

#### Stage 1 (shared)

Before any specific strategy is applied, we compute the minimal relaxation of the windows in Eq. (10) required to keep the problem feasible,

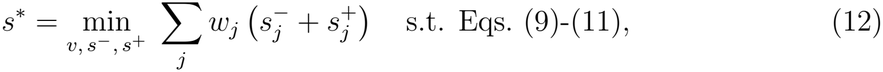

with weights

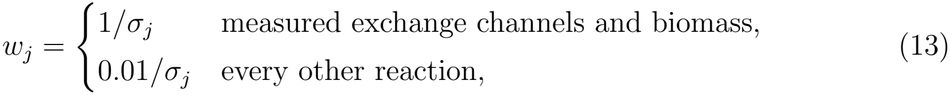

so that relaxing an intracellular bound is two orders of magnitude cheaper than relaxing a measured channel.

#### Stage 2 (strategy-specific)

Each strategy then optimises its own objective inside the relaxed polytope, with the total slack budget from Stage 1 frozen to within a small tolerance,

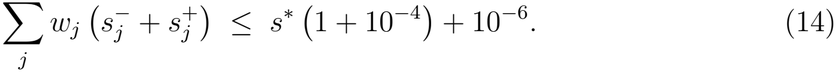

This two-stage construction fixes a single feasible set shared by all three strategies, so that any differences in the resulting reference fluxes are attributable to the choice of objective and not to the constraints.

#### Parsimonious FBA (pFBA)

In the pFBA strategy, the squared Euclidean norm of the flux vector is minimised:

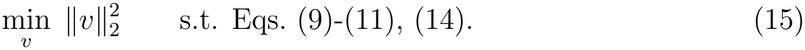

We note that Eq. (15) is a squared *L*_2_ objective rather than the sum of absolute fluxes (*L*_1_) used in the classical formulation of parsimonious FBA; we retain the name pFBA for consistency.

#### Max ATP

In the max ATP strategy, the flux through the reactions of the network that produce ATP is maximised and regularised by a small quadratic term to select a unique optimum among the flux distributions that achieve the same ATP yield:

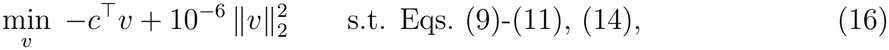

where *c*_*j*_ = 1 for the reaction(s) designated as the network’s ATP objective and *c*_*j*_ = 0 otherwise. The objective reactions are network-specific and are presented in Table 4.

#### Flux sampling

In flux sampling, the solution space is sampled, via constrained hit-and-run sampling. The result is a distribution of feasible fluxes. This proceeds in three steps at each time point.

#### Walk box

The Stage-1 relaxed windows are widened by max(0.01 × width, 10^−6^) and intersected with the physiological box in Eq. (11). This widening is necessary because the Stage-1 solution (Eq. (12)) generically sits exactly on the relaxed window boundary, which would otherwise pin the sampler’s starting point to a vertex.

#### Reachable range and interior point

Flux variability analysis within the walk box gives a per-reaction range [*e*_*lb,j*_, *e*_*ub,j*_], with half-width *d*_*j*_ = (*e*_*ub,j*_ − *e*_*lb,j*_)*/*2. A per-reaction-scaled Chebyshev centre of the walk box is used to seed the sampler,

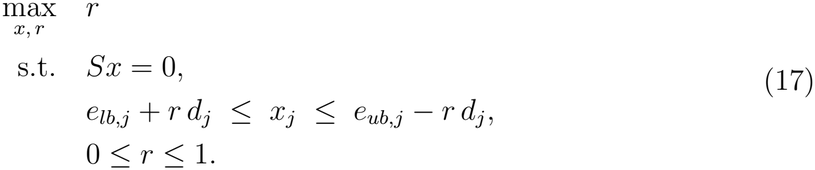

At *r* = 1 every reaction sits at the midpoint of its reachable range, regardless of the range’s width.

#### Hit-and-run sampling

Starting from the interior point of Eq. (17), samples are drawn by constrained hit-and-run sampling confined to null(*S*), with step directions scaled by each reaction’s half-width *d*_*j*_ so that the walk mixes at a comparable relative rate across reactions of very different flux magnitude. Every tenth step is retained, for a default of *n*_samples_ = 200 draws per time point. The resulting sample mean 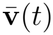 is projected back onto the exact relaxed windows via a small quadratic program in scaled coordinates, so that it remains feasible with respect to Eqs. (9)-(10); the sample standard deviation per reaction is denoted *σ*_ref_ (*t*).

#### Incorporation of the reference flux into training

Each reference strategy enters the training loss through a reference term ℒ_ref_, restricted to reactions that are not already constrained directly by data. For the point-valued strategies (pFBA, max ATP),

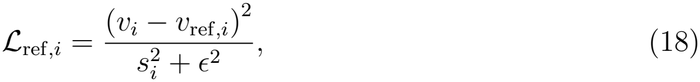

while for the sampling strategy,

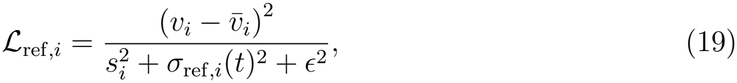

where *s*_*i*_ is the time point standard deviation of the reference flux for reaction *i*. Comparing Eqs. (18) and (19), the sampling form is equivalent to the point form rescaled by 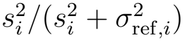, which down-weights reactions that the feasible polytope leaves comparatively undetermined.

### Error Calculation

The normalised MAE is calculated as follows:

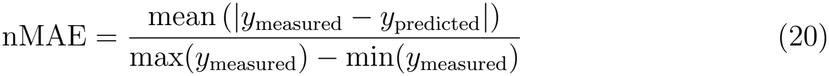

A normalised MAE of 0.2 means the average absolute error is 20% of the measured variable range for that metabolite in that held-out experiment.

### Part II: Model Performance

### Part II: Network Size

### Part II: Optimisation Strategy

**Table 7:**
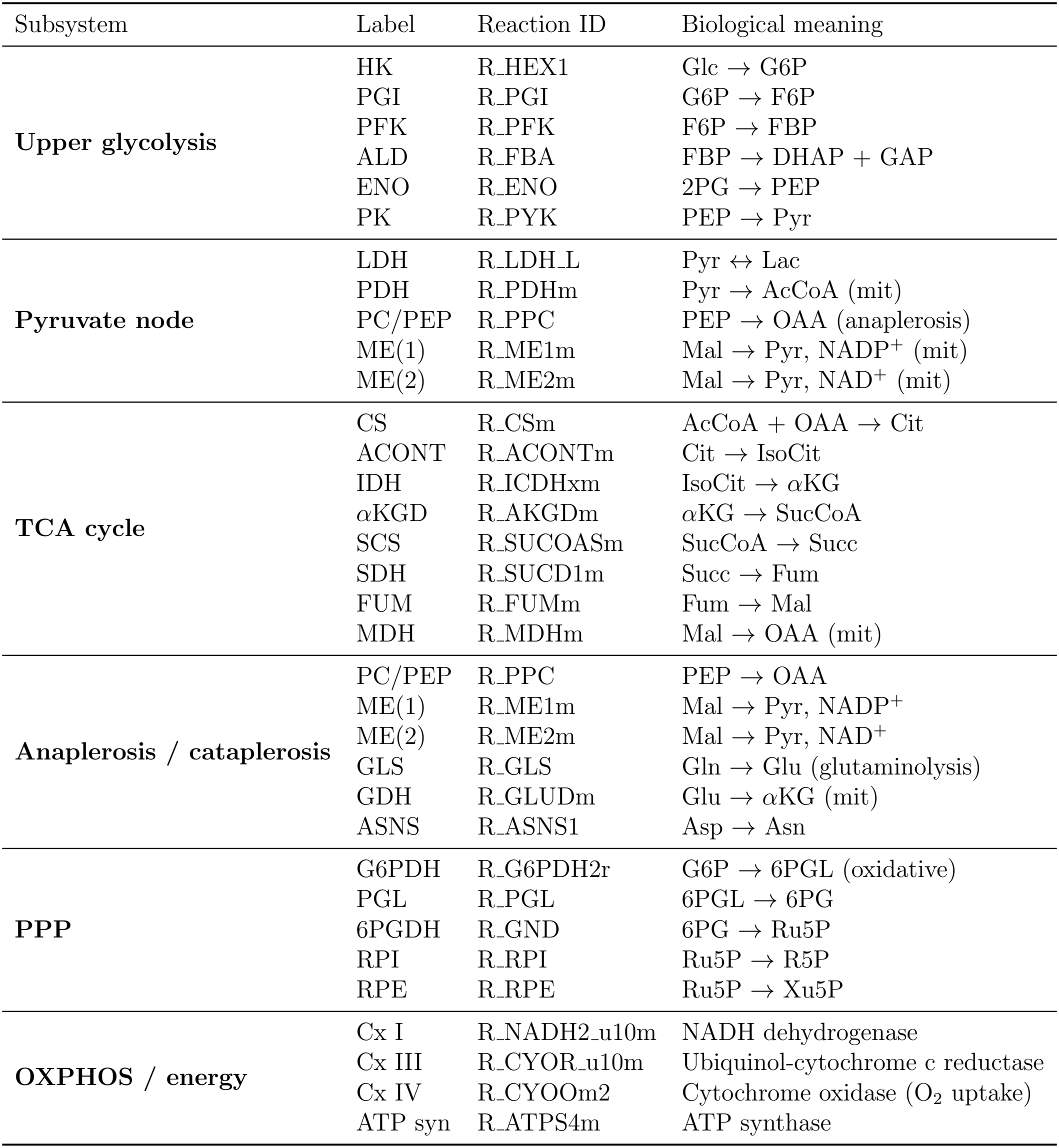
iCHO2441 reaction map (R IDs).

| Subsystem | Label | Reaction ID | Biological meaning |
|---|---|---|---|
| **Upper glycolysis** | HK | R_HEX1 | Glc → G6P |
|  | PGI | R_PGI | G6P → F6P |
|  | PFK | R_PFK | F6P → FBP |
|  | ALD | R_FBA | FBP → DHAP + GAP |
|  | ENO | R_ENO | 2PG → PEP |
|  | PK | R_PYK | PEP → Pyr |
| **Pyruvate node** | LDH | R_LDH_L | Pyr ↔ Lac |
|  | PDH | R_PDHm | Pyr → AcCoA (mit) |
|  | PC/PEP | R_PPC | PEP → OAA (anaplerosis) |
|  | ME(1) | R_ME1m | Mal → Pyr, NADP<sup>+</sup> (mit) |
|  | ME(2) | R_ME2m | Mal → Pyr, NAD<sup>+</sup> (mit) |
| **TCA cycle** | CS | R_CSm | AcCoA + OAA → Cit |
|  | ACONT | R_ACONTm | Cit → IsoCit |
|  | IDH | R_ICDHxm | IsoCit → αKG |
|  | αKGD | R_AKGDm | αKG → SucCoA |
|  | SCS | R_SUCOASm | SucCoA → Succ |
|  | SDH | R_SUCD1m | Succ → Fum |
|  | FUM | R_FUMm | Fum → Mal |
|  | MDH | R_MDHm | Mal → OAA (mit) |
| **Anaplerosis / cataplerosis** | PC/PEP | R_PPC | PEP → OAA |
|  | ME(1) | R_ME1m | Mal → Pyr, NADP<sup>+</sup> |
|  | ME(2) | R_ME2m | Mal → Pyr, NAD<sup>+</sup> |
|  | GLS | R_GLS | Gln → Glu (glutaminolysis) |
|  | GDH | R_GLUDm | Glu → αKG (mit) |
|  | ASNS | R_ASNS1 | Asp → Asn |
| **PPP** | G6PDH | R_G6PDH2r | G6P → 6PGL (oxidative) |
|  | PGL | R_PGL | 6PGL → 6PG |
|  | 6PGDH | R_GND | 6PG → Ru5P |
|  | RPI | R_RPI | Ru5P → R5P |
|  | RPE | R_RPE | Ru5P → Xu5P |
| **OXPHOS / energy** | Cx I | R_NADH2_u10m | NADH dehydrogenase |
|  | Cx III | R_CYOR_u10m | Ubiquinol-cytochrome c reductase |
|  | Cx IV | R_CYOOm2 | Cytochrome oxidase (O<sub>2</sub> uptake) |
|  | ATP syn | R_ATPS4m | ATP synthase |

### Part II: Regime-specific Adaptations

**Figure 19:**
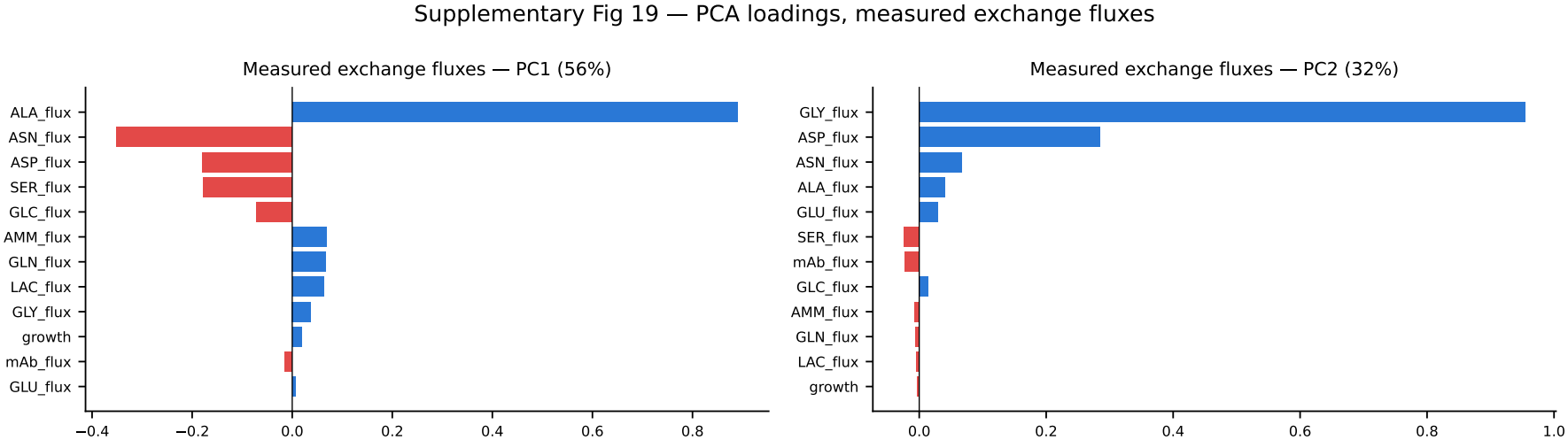
PCA loadings for exchange fluxes.

## Notes

### Competing Interest Statement

The studentship of M.M. was funded by BBSRC and GSK. S.B., R.T., G.N. and P.K. are employees of GSK.

### Summary of Updates

Manuscript revised for clarity; Table 2 corrected; Supplementary Information updated.

### Funding

- BBSRC
- GSK

